# Strigolactones are involved in enhancing iron uptake in maize

**DOI:** 10.1101/2024.10.29.620872

**Authors:** Stavroula Fili, Jiahn-Chou Guan, Karen E. Koch, Elsbeth L. Walker

**Affiliations:** Biology Department, University of Massachusetts, Amherst; Plant Biology Graduate Program, University of Massachusetts, Amherst; Horticultural Sciences Department, University of Florida, Gainesville, Florida 32611, U.S.A; Plant Molecular Biology Program, University of Florida, Gainesville. Florida 32611, U.S.A; Genetics Institute, University of Florida, Gainesville, Florida, 32610, U.S.A

## Abstract

Strigolactones are plant hormones with roles in a wide range of signaling and developmental processes. A yellow-striped maize mutant, (*inter<u>v</u>einal <u>y</u>ellow*) *ivy*, was determined to have low iron in tissues under normal growth conditions. The gene underlying the *ivy* mutation was mapped and identified as *ZmCCD8*, a key enzyme in the biosynthesis of strigolactones. Under iron-replete conditions, comparison of the transcriptomes of wild-type plants and maize *ccd8* mutants revealed suppression of several iron-regulated genes in *ccd8*. These genes are normally up-regulated during iron deficiency and include the key iron-regulated transcription factor *IRO2* as well as genes involved in the biosynthesis of iron chelators and transporters. External supply of synthetic strigolactone to *ivy* mutants alleviated chlorosis and returned iron-regulated gene expression to wild-type levels. In iron limited conditions, iron-regulated gene expression in *ccd8* mutants responded normally, indicating that strigolactones are not required for response to externally imposed iron deficiency. However, they are required for basal expression of iron-regulated genes when adequate iron is available, highlighting a distinction between iron homeostasis during normal growth, and the iron deficiency response triggered by the lack of external available iron. The connection between strigolactones and iron homeostasis is not limited to maize, as Arabidopsis *ccd8* mutants also show strong chlorosis when grown on medium with moderate levels of iron. This previously unappreciated role may have implications for the use of strigolactones in agricultural contexts.

## Introduction

Iron is an essential micronutrient for plant growth and development that is required for chlorophyll biosynthesis and cellular respiration, among many other processes. Iron deficiency in plants can lead to stunted development and poor yield. Humans also need iron for cellular respiration and hemoglobin biosynthesis; insufficient iron intake can lead to iron deficiency, which is the cause of 50% of human anemia cases worldwide (Turawa et al., 2021). Thus it is imperative that adequate iron is available in human diets. Although iron is abundant in soil, its low solubility makes it poorly bioavailable; thus, plants have developed mechanisms to increase its primary uptake (Marschner and Römheld, 1994). Most plants use an acidification-reduction strategy to acquire iron from the soil (Strategy I), while plants belonging to the grass family (*Poaceae*) have evolved a distinct strategy (Strategy II) based on the secretion of iron-chelating molecules called phytosiderophores (PS). Grasses include most of the world’s staple crops such as maize and rice.

Maize mutants have been essential in identifying core components of the Strategy II (grass) uptake system. The classic *yellow stripe 1* (*ys1*) (von Wiren et al., 1994; Curie et al., 2001) and *yellow stripe 3* (*ys3*) mutants (Nozoye et al., 2013; Chan-Rodriguez and Walker, 2018) helped to define the transporters for secretion of PS from root cells and subsequent uptake of PS-Fe complexes from the rhizosphere. *YS1* encodes the root iron-PS uptake transporter YS1(Curie et al., 2001), while *YS3* encodes the PS exporter TOM1(Chan-Rodriguez and Walker, 2018). These mutants show typical iron-deficiency chlorosis (yellow regions between green veins), which can be alleviated with foliar iron application. A recent study identified the *maize interveinal chlorosis 1* (*mic1*) mutant with a similar phenotype to *ys1* and *ys3* (Sun et al., 2022). *Mic1* encodes a 5′- methylthioadenosine nucleosidase (MTN) involved in nicotianamine (NA) biosynthesis. Nicotianamine is an internal iron chelator, and is also the precursor for PS biosynthesis. The *mic1* mutants are partially rescued by iron supplementation only if combined with exogenous NA application (Sun et al., 2022).

To identify new genes involved in iron homeostasis in maize, we screened publicly available maize mutants for iron-deficiency (yellow striped) phenotypes and tested whether they can be reversed with foliar iron supplementation (Chan-Rodriguez and Walker, 2018). During this process, we identified a new yellow-striped mutant which we named *inter<u>v</u>einal <u>y</u>ellow* (*ivy*;(Chan-Rodriguez and Walker, 2018). Leaves of *ivy* plants grown in soil in the greenhouse contained significantly low iron levels. Complementation was observed following crossing to both *ys1* and *ys3* mutants, indicating that *IVY* is another maize gene that is involved in iron homeostasis in maize (Chan-Rodriguez and Walker, 2018).

Strigolactones (SLs) were first identified due to their ability to stimulate the germination of seeds of *Striga* parasitic plants (Cook et al., 1966; Siame et al., 1993), but were later recognized as having important roles in the determination of plant form (Gomez-Roldan et al., 2008; Umehara et al., 2008). The SL-related mutants were first identified as excessive branching and dwarf mutants and were named MAX (more axillary growth) in Arabidopsis (Sorefan et al., 2003; Booker et al., 2004; Booker et al., 2005; Waters et al., 2012a; Waters et al., 2012b; Li et al., 2022), and *D*/*HTD* (*dwarf*/*high tillering dwarf*) in rice(Zou et al., 2006; Arite et al., 2007; Arite et al., 2009; Jiang et al., 2013; Zhang et al., 2014). SL hormones are derived from the carotenoid pathway (Matusova et al., 2005). Mutants in genes encoding the SL biosynthetic enzymes, which include *D27* (β-carotene isomerase), *MAX3*/*CCD7* (*Carotenoid Cleavage Dioxygenase 7*), *MAX4*/*CCD8* (*Carotenoid Cleavage Dioxygenase 8*) and *MAX1* (cytochrome P450, CYP450), are not able to produce SL hormones, resulting in the characteristic branching and dwarf phenotype. Meanwhile, mutants in downstream signaling components, including *D14*, *D3*/*MAX2* and *D53*/*SMXLs* (*Suppressor of MAX2 Like*), produce SLs but cannot perceive them, which results in a similar SL-deficient phenotype. Use of these mutants led to our current understanding that SLs bind to the D14 receptor (Hamiaux et al., 2012), which recruits the F-box protein D3/MAX2 and leads to the degradation of D53/SMXLs(Jiang et al., 2013; Zhou et al., 2013a; Yao et al., 2018; Li et al., 2022). D53 is a transcriptional repressor of SL signaling (Jiang et al., 2013), so upon SL perceived by D14, D53 repression is released, and the expression of downstream targets is induced (Waters et al., 2017; Liu et al., 2021; Guan et al., 2023).

Here we report that we have mapped the underlying mutation in *ivy* and found the responsible gene, *CCD8*, which is critical for the production of the SL family of plant hormones. A connection between SLs and iron homeostasis has not been previously reported, although Guan et al. (Guan et al., 2023) recently described the yellow striped leaf phenotype of the SL-deficient *ccd8* mutant. Here, we show that the iron deficiency phenotype is a result of the absence of SLs in these mutants. We also show that the mutant has lower than normal expression of iron-responding genes, and that this low expression can be reversed to wild-type levels by external SL application. A mutant in CCD7, an enzyme catalyzing another step in SL biosynthesis, showed a similar, yellow-striped phenotype. SL biosynthetic mutants in both rice and Arabidopsis also exhibit yellow striped phenotypes. Our results support the conclusion that SLs are involved in the regulation of iron homeostasis in maize.

## Results

### The *ivy* mutant is a monogenic recessive and has an iron-deficiency phenotype

The *ivy* mutant seed stock was obtained from the Maize Genetics Cooperation Stock Center (MGCSC #3812O; *ys*^∗^-*N2398*), which listed it as a *yellow stripe* mutant with a recessive pattern of inheritance, generated by EMS mutagenesis (Chan-Rodriguez and Walker, 2018). Individuals from the *ivy* Maize COOP stock were self-pollinated, and their progeny were crossed to inbred lines W22 and B73 to produce segregating populations. The F2 populations segregated *ivy* and wild-type individuals in a 1:3 phenotypic ratio, demonstrating a monogenic recessive mutation. In order to confirm the low iron phenotype described previously for soil-grown plants (Chan-Rodriguez and Walker, 2018), we grew an F2 family that came from the cross to W22 in hydroponic culture. The iron concentration in leaves and roots of mutant (yellow striped) and WT siblings from this segregating population was determined using ICP-MS. The leaves of *ivy* mutant plants had low iron compared to their wild-type siblings (Figure 1A). There was no significant difference in the iron concentration in the roots of mutant and WT siblings (Figure 1B).

**Figure 1:**
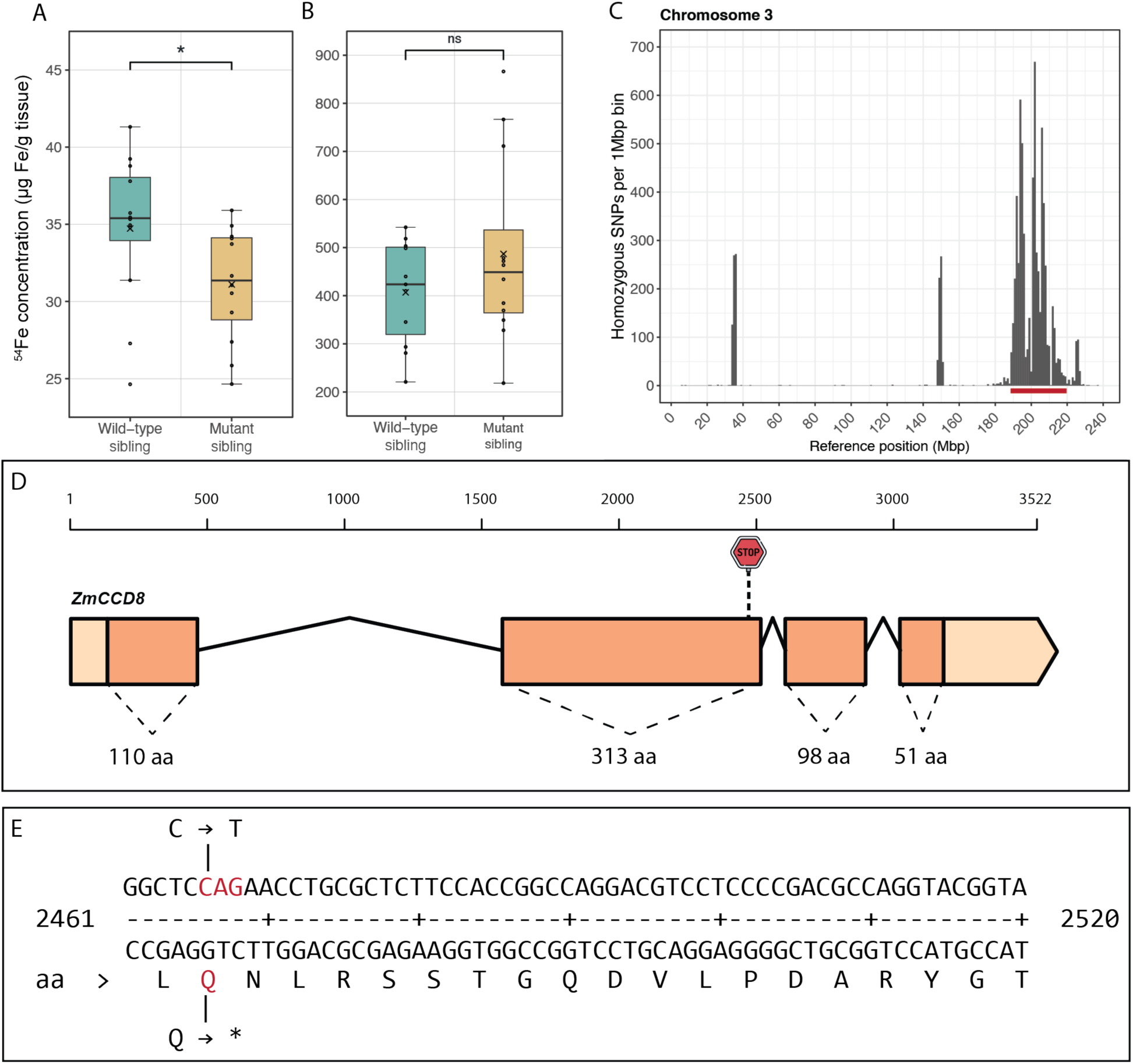
The iron-deficient *ivy* mutant. (A, B) Iron concentration of *ivy* and wild-type siblings growing in hydroponic culture. Measurements were performed by ICP-MS for whole shoots (A) and whole roots (B). Significance was determined using the Mann – Whitney test (n=12). Asterisks (*) indicate significance (* p < 0.05, ** p < 0.01, *** p < 0.001, **** p < 0.0001). (C) Plot of homozygous variants on chromosome 3, for *ivy* mutant pool plants compared to the W22 reference genome. (D) Representation of the *CCD8* gene in maize and the relative location of the stop codon introduced in *ivy* (exon 2). (E) Exact location of the nonsense mutation present in the *ivy* mutant.

### The *ivy* gene maps to the *CCD8* locus

The F2 population generated by crossing to W22 was used as a mapping population, and bulked-segregant analysis coupled with next-generation sequencing (BSA-seq) was employed to identify the underlying mutation. Reads from individually sequenced mutant and wild-type pools were aligned to the W22 reference genome (assembly version: *Zm-W22-REFERENCE-NRGENE-2.0*) to reveal homozygous variants present in the mutant but not in the wild-type siblings (Supplemental Figure S1). Multiple regions of homozygosity were present on several chromosomes. Previous mapping of another mutant crossed to the same W22 stock (unpublished), revealed that all but one of these regions were also called as homozygous in that analysis. Thus, we were able to ignore these regions, which aided in selection of a single region of homozygosity associated specifically with the *ivy* mutation, on the long arm of chromosome 3 (Figure 1C). We performed SNP filtering in the region between 188,000,000 and 218,000,000 bp to exclude any non-EMS type mutations, since the *ivy* stock was generated by EMS mutagenesis. Then, we identified variants that occur in eight sequenced maize lines (Hirsch et al., 2016; Jiao et al., 2017; Springer et al., 2018; Sun et al., 2018; Haberer et al., 2020; Wang et al., 2023) and excluded those, as they are unlikely to be responsible for the *ivy* phenotype when homozygous. Finally, we analyzed the effect of the remaining SNPs at the protein level and selected those that had either a moderate or high effect predicted on the encoded protein as candidates.

Out of 32 candidate genes meeting all the above criteria, only 2 had a predicted high effect on the encoded protein. One of these genes (*Zm00004b018959* in the W22 annotation; Zm00001eb153000 in B73v5) encodes the carotenoid cleavage dioxygenase, CCD8, which is a key enzyme in the biosynthesis of SLs (Sorefan et al., 2003; Gomez-Roldan et al., 2008; Umehara et al., 2008) (Figure 1D). Sequence comparison between the mutant and wild-type DNA pools revealed a single nucleotide C to T substitution in codon 408 of the *CCD8* CDS (Figure 1E). This mutation causes a premature stop codon (Gln408*/572), resulting in a protein product that is 164 amino acids shorter than wild-type. The mutation was confirmed by sequencing the relevant region in homozygous progeny from the MGCSC stock.

To test whether a mutation in *CCD8* is responsible for the *ivy* phenotype, we performed complementation tests with a previously characterized mutant allele of *CCD8*. This mutant allele, *ccd8::Ds*(*zmccd8*) has a *Ds6*-*like* insertion at the junction of the second intron and third exon (Guan et al., 2012). *zmccd8* plants grown in the field next to *ivy* plants showed the same yellow striped phenotype as *ivy* mutant plants and are similar to the *ys1* mutant (Figure 2A). Both *zmccd8* and *ivy* mutant plants had short stature relative to W22 or WT siblings in segregating populations. We obtained three independent complementation crosses between *ivy* and *zmccd8* mutants. All progeny resulting from these three crosses presented the typical yellow striped phenotype observed in both parents, confirming that *ivy* is an allele of *ccd8* (Figure 2B and C).

**Figure 2:**
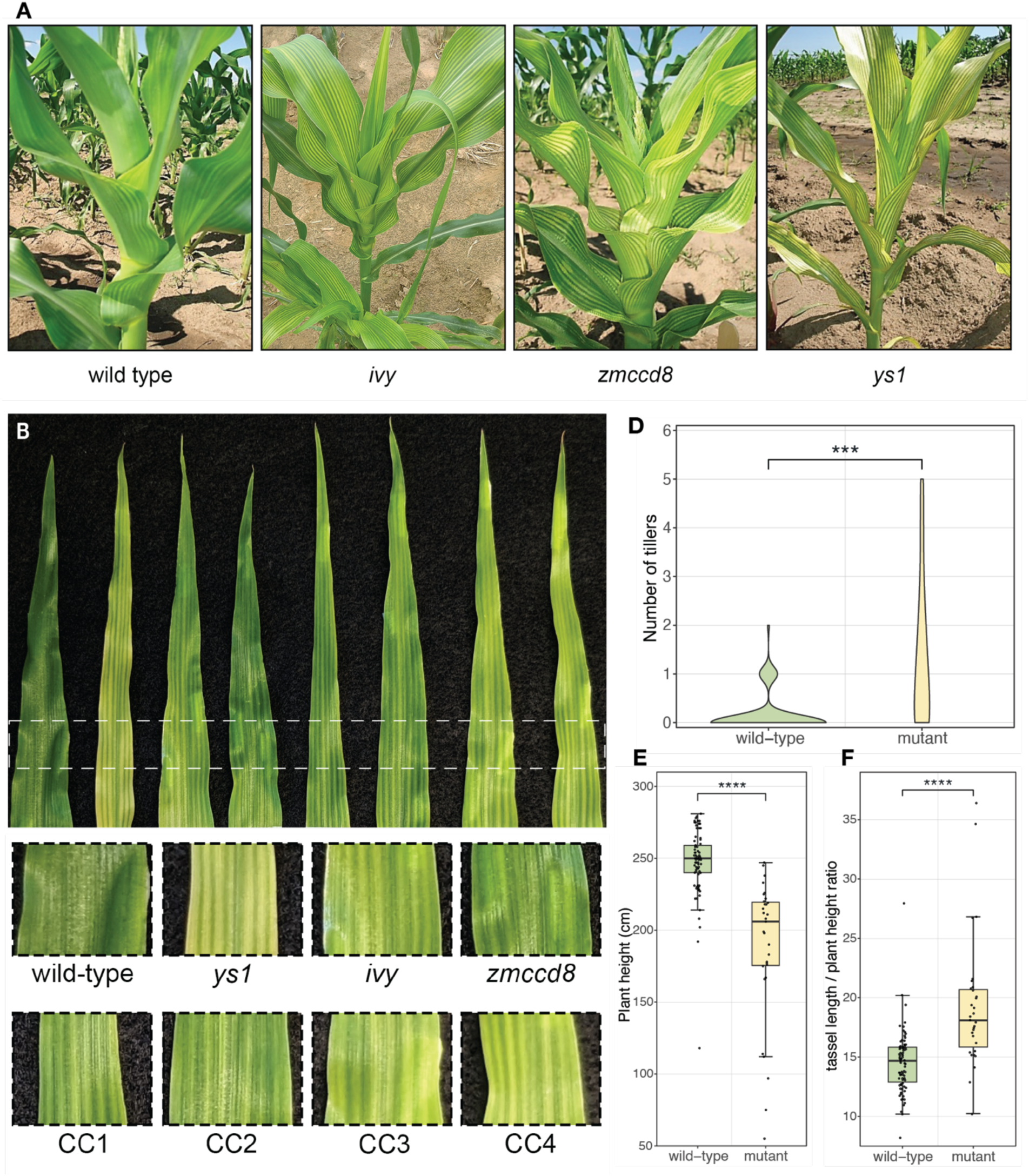
Phenotype and non-complementation of *ivy* and *ccd8* mutants. (A) Yellow stripe phenotypes of *ivy* and *ccd8* compared to *ys1* mutant and wild-type. (B, C) Results of complementation crosses between *ivy* and *zmccd8* mutants. Leaves from F1 progeny (CC1, CC2, CC3 and CC4) compared to *ys1*, *ivy* and *zmccd8* homozygotes. Plants were grown in soil in the greenhouse and pictures were taken 19 days after sowing. (D, E, F) Morphological measurements in *ivy* mutants compared to wild-type siblings, from an F2 population generated by crossing *ivy* to W22. Significance was determined using Student’s *t*–test (unpaired, unequal variance). Asterisks (*) indicate significance (* p < 0.05, ** p < 0.01, *** p < 0.001, **** p < 0.0001).

Previous characterization of the *zmccd8* mutant revealed a variety of developmental phenotypes related to SL hormone signaling, including increased branching, shorter plant height, and drooping tassels (Guan et al., 2012). We examined these traits in the *ivy* mutant by observing mutant individuals segregating from the two populations (W22 and B73) mentioned earlier. In both populations, mutant siblings showed an increase in the number of branches (tillers) compared to wild-type siblings (Figure 2D; Supplemental Figure S2A). Mutant siblings were also typically shorter, and the vast majority presented the drooping tassel phenotype, resulting from a slender stem (rachis) and an increased tassel height to plant height ratio (Figure 2E and F; Supplemental Figure S2B and C).

### The *ivy* phenotype is caused by a lack of SLs and not by a non-SL phenomenon

CCD8 is expected to act enzymatically on β-apo-10’-carotenal to produce carlactone (Figure 3A), but it is possible that it might have previously unappreciated enzymatic activity outside of the SL biosynthetic pathway. If that were the case, then the iron-deficient phenotype of maize *ccd8* mutants could result from this other activity. We sought to understand whether the iron deficiency symptoms of the *ccd8* mutant are caused by a lack of SL *per se* by testing whether the synthetic SL analog, *rac*-GR24, is capable of reversing the chlorosis of *ccd8* mutant plants. External supply of *rac*-GR24 in SL mutants has been previously shown to revert their developmental phenotypes (Gomez-Roldan et al., 2008; Umehara et al., 2008; Ruyter-Spira et al., 2011). We grew *ivy* and wild-type W22 plants in hydroponic culture with normal iron nutrition for a period of ten days. Throughout this period one set of the *ivy* plants was provided with *rac*- GR24 in the culture medium, while the control *ivy* plants and the W22 control plants did not receive *rac*-GR24.

**Figure 3:**
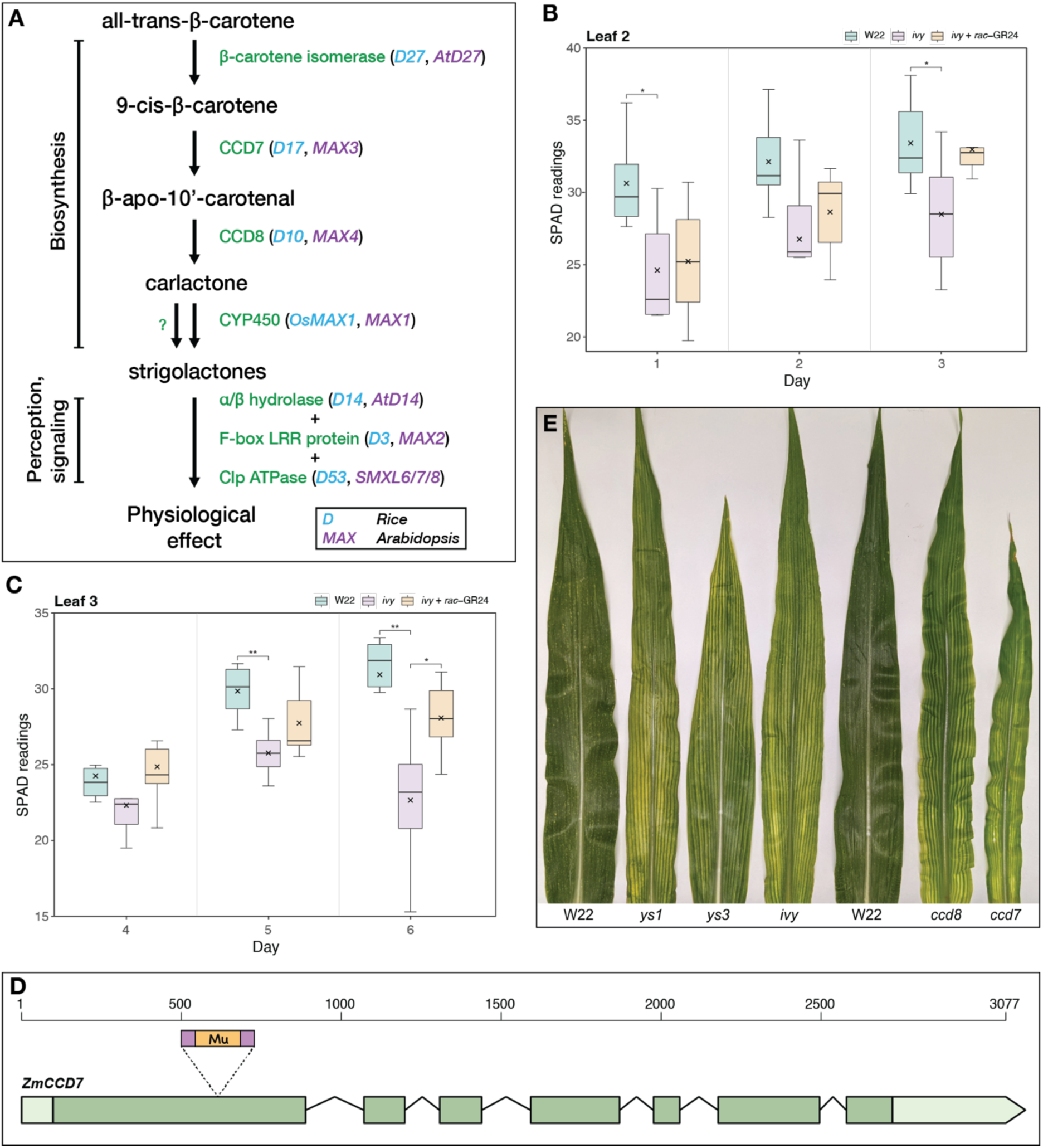
Interveinal chlorosis of *ivy* mutants is alleviated by external *rac*-GR24 application. (A) The strigolactone biosynthetic pathway. The diagram shows known genes involved in the biosynthesis and the perception of strigolactones. The gene names are given in the rice (blue) and Arabidopsis (purple) nomenclature, according to (Booker et al., 2005; Zou et al., 2006; Arite et al., 2007; Arite et al., 2009; Nelson et al., 2011; Alder et al., 2012; Zhang et al., 2014; Abuauf et al., 2018; Wang et al., 2020a). The gene product is given in green. (B, C) Chlorosis (SPAD values) was assessed in a 3-day window for the second (B) and third (C) leaves. Statistical analysis was performed using one-way ANOVA with posthoc Tukey HSD (n=6). Asterisks (*) indicate significance (* p < 0.05, ** p < 0.01, *** p < 0.001, **** p < 0.0001). (D) Representation of the structure of *CCD7* gene in maize and relative location of the insertion (exon 1) present in the *ccd7* mutant (stock: mu-illumina_578516.6). (E) Comparison of the different yellow-striped mutants grown in the field. The flag leaf is shown for each genotype. The iron deficient *ys1* and *ys3* mutants are shown for reference. *ivy* is the mutant identified in this study, *ccd8* is the *ccd8::Ds* and *ccd7* is a *Mutator* insertion mutant (stock: mu-illumina_578516.6) (Williams-Carrier et al., 2010).

First, the efficacy of the *rac*-GR24 treatment was assessed by quantifying the emergence of crown roots in treated and untreated plants. Crown root emergence is delayed in maize *ccd8* mutants (Guan et al., 2012). Untreated *ivy* plants showed delayed emergence of the crown roots compared to wild-type plants (Supplemental figure S3). However, crown root emergence in *ivy* plants treated with *rac*-GR24 was almost identical to wild-type plants, thus confirming the effectiveness of the hormone treatment (Supplemental figure S3). Having established that the *rac*-GR24 treatment was effective, we examined chlorosis visually and by measuring the relative chlorophyll amount (Gianquinto et al., 2004; Hawkins et al., 2007). *ivy* plants supplemented with *rac*-GR24 showed higher chlorophyll levels than untreated *ivy* plants, which had significantly lower values than wild-type controls (Figure 3B and C). Because supplementation with SL is able to reverse chlorosis, we conclude that the lack of SLs in the *ivy* mutant is responsible for the yellow-stripe phenotype. This result suggests that SLs directly influence iron homeostasis in maize.

To further test the hypothesis that SLs are involved in iron homeostasis, we examined whether other SL-related mutants in maize exhibit iron-deficiency phenotypes. We obtained mutant stocks for another gene from the SL pathway: *CCD7.* CCD7 is a carotenoid cleavage dioxygenase involved in SL biosynthesis that acts upstream of CCD8 (Booker et al., 2004; Pan et al., 2016) (Figure 3A). Maize *CCD7* is able to complement Arabidopsis *max3* mutants, demonstrating that it is a *bona fide* SL biosynthetic gene (Pan et al., 2016). A transposon insertion mutant for *ccd7* was obtained from the Mu-Illumina project (mu-illumina_578516.6, Zm00001eb074640) (Williams-Carrier et al., 2010). The stock segregated yellow striped individuals. These individuals were confirmed to be homozygous for a *Mutator* (*Mu*) insertion in the first exon of the *CCD7* gene (Figure 3D). The *ccd7* mutants were grown in the field alongside *ivy* and *ccd8::Ds* mutants for direct comparison. Homozygous *ccd7* mutants started to show yellow striping during the early stages of their development and this continued throughout their life cycle, similar to *ivy* and *ccd8* mutants (Figure 3E). These results support the idea that the lack of SLs in maize mutants is responsible for the observed yellow-stripe phenotype.

### The iron deficiency phenotype in *ivy* is unlikely to be caused by developmental alterations of the roots

Having established that the iron deficiency phenotype in *ivy* is a consequence of SL deficiency, we hypothesized that the delayed emergence of crown roots in *ivy* mutants could cause the plants to be anatomically impaired in iron uptake. Crown root development is stimulated by SL (Arite et al., 2012), and *ccd8* mutant plants have reduced crown root development (Guan et al., 2012). The idea of anatomical impairment of iron uptake was strengthened by the observation that key genes involved in iron homeostasis (*ZmYS1*, *ZmTOM1*, *ZmOPT3*, *ZmDMAS1*, *ZmNAAT1*, *ZmNAS1;1*, *ZmNAS1;2*, *ZmNAS2;1*, *ZmNAS2;2*, *ZmNAS6;1*, *ZmNAS6;2*, *ZmIRO2;1*, *ZmIRO2;2*, *ZmIRO3*, *ZmNRAMP1*). all showed higher expression in the crown roots compared to other root types ((Stelpflug et al., 2016); Supplemental figure S4A). This observation suggested that crown roots might be particularly important for iron uptake and that their delayed emergence in *ivy* could have a negative effect on iron uptake from the soil.

To address this question, we examined maize mutants showing defects in crown root development. We obtained seed stocks for *rootless concerning crown and seminal roots* (*rtcs*) and *rootless1* (*rt1)*, two mutants that are impaired in crown root development. *rtcs* has a complete absence of nodal roots (crown or brace roots) (Hetz et al., 1996; Taramino et al., 2007). *rt1* has a reduced number of crown roots, and it does not produce any later types of nodal roots (Hochholdinger et al., 2004). Neither of these mutants were previously reported to have a yellow-striped phenotype. We grew *rt1* and *rtcs* segregating stocks and observed the mutant individuals. Both mutants had delayed growth compared to their wild-type siblings; however, neither of these mutants exhibited a chlorotic phenotype at any stage of their development (Supplemental figure S4B and C). We conclude that the delayed emergence of crown roots in *ivy* is unlikely to be the cause of chlorosis observed in *ivy* mutants. While crown roots may be important for iron uptake, their absence is apparently compensated by other types of roots. We conclude that the yellow-striped phenotype observed in *ivy* most likely results from another aspect of SL hormonal action.

### Transcriptomic analysis of maize roots and shoots under normal, deficient, and resupplied iron conditions

As part of our search for novel genes involved in iron homeostasis in maize, we performed a transcriptomic study to identify genes that respond to iron deficiency. Maize W22 plants were grown in hydroponic culture with sufficient (+Fe), deficient (−Fe), and resupplied (±Fe) iron conditions. All plants were initially grown in iron-sufficient solution; then −Fe and ±Fe plants were grown in iron-deficient solution for three days. After the deficiency period, the ±Fe plants were resupplied with iron-sufficient solution for 24 hours, while the −Fe plants received fresh iron-deficient solution. The resupplied iron condition was used to distinguish those genes that respond quickly to changes in iron availability in the environment.

A total of 216 genes in the roots and 71 genes in the shoots were differentially expressed between −Fe and +Fe plants with significant criteria (|FC| ≥ 2, P-adj (FDR) ≤ 0.01). In roots, 188 transcripts increased in abundance, and 28 transcripts decreased in abundance during iron deficiency. In shoots, 15 transcripts increased in abundance, and 56 transcripts decreased in abundance during iron deficiency.

Having established this set of iron-responding transcripts in maize (Supplemental data 1), we examined their behavior during iron resupply. 178 transcripts in roots and 63 transcripts in shoots returned to basal levels of expression or partially returned towards basal levels of expression after 24 hours of iron resupply. These are defined as “fast responding” genes, and we consider these likely to be directly involved in iron homeostasis during iron deficiency. Among these genes are *YS1* (Zm00001eb249020), *TOM1* (Zm00001eb133440), *NAS1* (Zm00001eb396230), *NAS2* (Zm00001eb014700), *NAS6* (Zm00001eb396110),NAS10 (Zm00001eb396280), *NAAT1* (Zm00001eb203230)*, DMAS1* (Zm00001eb010040) and homologs of *AtNRAMPs* (Zm00001eb231210 and Zm00001eb400560) and *AtOPT3* (Zm00001eb057440). All of these genes have established roles in adaptation to iron deficiency (Curie et al., 2000; Curie et al., 2001; Nozoye et al., 2013; Zhou et al., 2013b; Mendoza-Cozatl et al., 2014; Zhai et al., 2014; Castaings et al., 2016; Peris-Peris et al., 2017; Chan-Rodriguez and Walker, 2018; Zhang et al., 2022). Smaller sets of transcripts (38 in roots and 8 in shoots) did not significantly shift towards basal expression levels during resupply. These were categorized as slow responding. Among these slow responding genes only two have previously characterized roles in the iron deficiency response. One is Zm00001eb121190, which may be involved methionine scavenging and therefore related to phytosiderophore biosynthesis. The other is *Nramp6* (Zm00001eb231210) which encodes a metal transporter.

An observation that came out of this RNA-seq analysis was the strong iron-deficiency response of Zm00001eb140680, annotated as *ZmbHLH126* in MaizeGDB and as *IRON-RELATED TRANSCRIPTION FACTOR 2* (*IRO2*) in NCBI. *OsIRO2* is an important transcription factor involved in iron homeostasis in rice (Mizuno et al., 2003; Ogo et al., 2006; Wang et al., 2020b), targeting key components of the iron uptake machinery, such as the iron-phytosiderophore transporter *OsYSL15* and the phytosiderophore exporter *OsTOM1* (Ogo et al., 2007; Ogo et al., 2011). We performed a BLAST search against the maize genome using the sequence of OsIRO2 as a query, and obtained two hits with significant similarities to OsIRO2: Zm00001eb140680 (here referred to as *ZmIRO2;1*) and Zm00001eb362800 (annotated as *ZmbHLH54* in MaizeGDB, here referred to as *ZmIRO2;2*) (Supplemental figure S5). Synteny analysis revealed conserved gene sequences between the region around *OsIRO2* (rice chromosome 1), and the regions around both *ZmIRO2;1* and *ZmIRO2;2* (maize chromosomes 3 and 8, respectively) (Supplemental figure S5). Maize chromosomes 3 and 8 are known to contain duplicated regions that correspond to rice chromosome 1 (Ahn and Tanksley, 1993; Salse et al., 2004). Thus, *OsIRO2* has two orthologs in maize. In our RNA-seq dataset, the expression of *ZmIRO2;2* was only mildly up-regulated in –Fe (FC=1.88, P-adj=3.23E-17 in roots, FC=1.99, P-adj=1.57E-36 in shoots) and thus did not pass our strict criteria as an iron-regulated gene. Another transcriptomic study using microarrays showed a 2-fold change for *ZmIRO2;2* in –Fe roots (Zanin et al., 2017). At this point, the respective roles of ZmIRO2;2 and ZmIRO2;1 in relation to iron homeostasis are unclear.

Notably, we found that genes encoding SL biosynthetic enzymes or those involved in SL signaling (*CCD8/*Zm00001eb153000; *D53*/Zm00001eb404750; *D14A*/Zm00001eb007670; *D14B*/Zm00001eb400720; *CCD7* /Zm00001eb074640; *ZmMax1b*/Zm00001eb123090; *ZmCLAMT1*/Zm00001eb123070; and *ZmCYP706C37*/Zm00001eb123110) were not among the iron regulated genes that we identified. This highlights that while SL modulates the level of iron-related genes, iron levels do not appear to modulate the levels of SL-associated genes, at least at the level of mRNA abundance during short term iron starvation.

### Iron-regulated gene expression is impaired in *ivy* mutants

To better understand the influence of SLs in iron homeostasis, we asked whether *ivy* plants have any changes in the expression of iron-regulated genes. We used RNA-seq to compare the transcriptomes of the SL-deficient *ccd8* mutant to those of wildtype (B73) using roots and shoots of 14-day-old seedlings. Differentially expressed genes (|FC| ≥ 2, P-adj (FDR) ≤ 0.01) were selected for further analysis. Overall, in *ccd8*, 99 genes in shoots and 151 genes in roots were up-regulated by 2-fold or more. 62 genes in shoots and 915 in roots were down-regulated (Supplemental data 2). Genes for iron homeostasis were among those significantly affected by SL deficiency. Two of the mRNAs down-regulated by more than 3-fold in roots are encoded by *ZmIRO2;1* and *ZmIRO2;2*. Also, down-regulation by SL deficiency was observed for most homologs of genes involved in biosynthesis of phytosiderophores, including precursors from the S-adenosyl-L-methionine cycle (Figure 4A). Also notable is that 7 out of 10 nicotianamine (NA) synthase (*ZmNAS1;1∼6;2*, Figure 4A) genes are down-regulated in the *ccd8* mutant roots and so too is *ZmDMAS1* (deoxymugineic acid synthetase; *Zm00001eb010040*), encoding the enzyme that converts NA to the phytosiderophore deoxymugineic acid (DMA). More modest (less than 2-fold) transcript reductions were significant for several more genes involved iron homeostasis (Supplemental data 2). Among these are *ZmYS1* (Zm00001eb249020), *ZmYSL3* (Zm00001eb133440), *ZmTOM2* (Zm00001eb196170), *ZmIRT1* (Zm00001eb052440), *ZmNRAMP1* (Zm00001eb400560), and *ZmMATE2*/*ZmPEZ1* (Zm00001eb219790). The expression change of *ZmIRO2;2*, *ZmYS1*, *ZmNAS6* and *ZmDMAS1* in *ccd8* mutants was confirmed by quantitative real-time RT-PCR analysis (Figure 4B).

**Figure 4:**
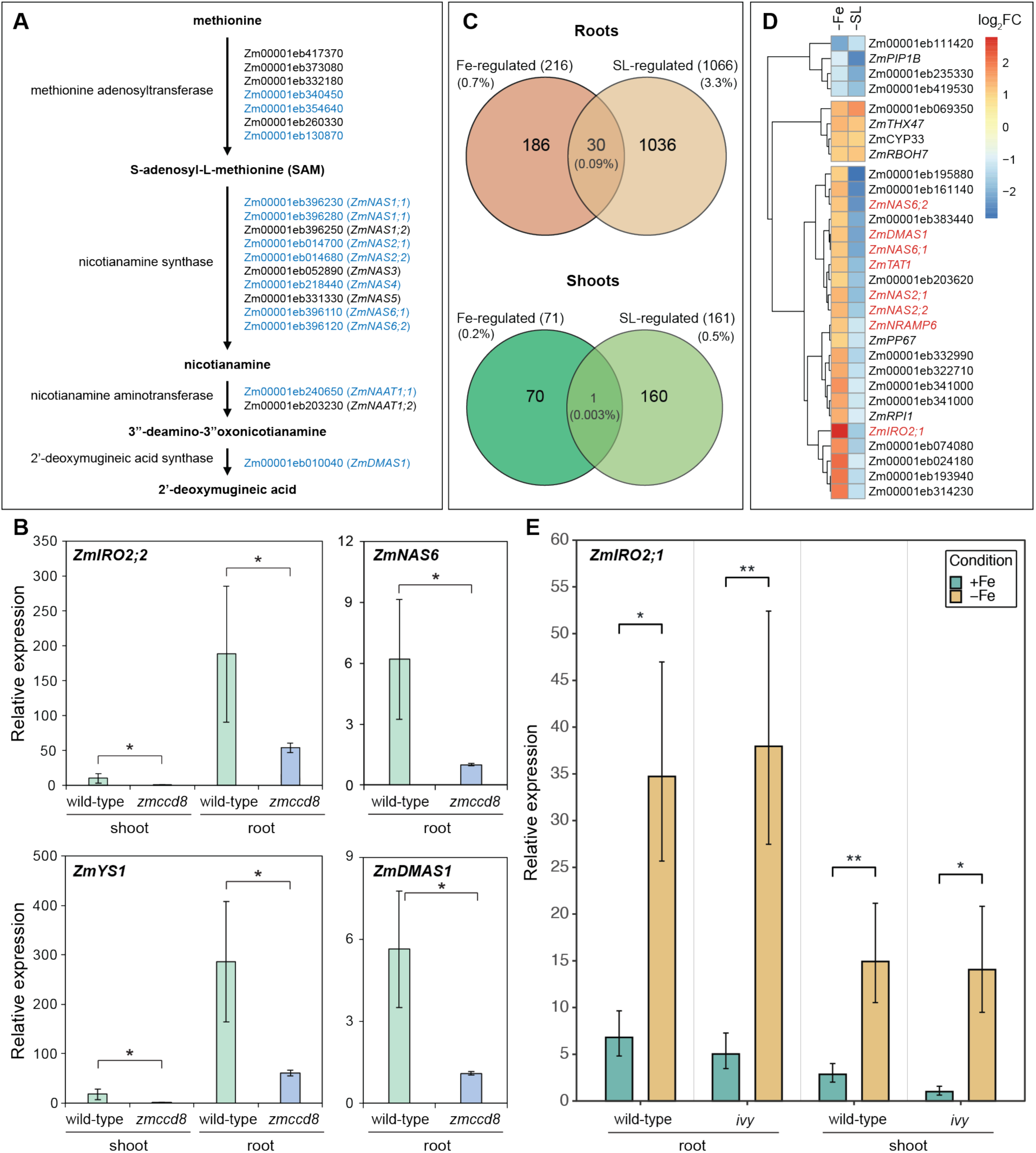
Iron-responsive gene expression in *ivy*/*ccd8*. (A) The pathway of phytosiderophore biosynthesis in maize, with enzymes and corresponding genes for each step shown in blue where down-regulated by 2-fold or more in roots. *NAS* gene nomenclature is according to Zhou et al (Zhou et al., 2013b). (B) Additional RT-qPCR analysis of differential expression for key genes from RNA-seq profiles: These included *ZmIRO2;2*, *ZmNAS6*, *ZmYS1*, or *ZmDMAS1*. Levels of *ZmIRO2;2* or *ZmYS1* mRNAs were evaluated in both shoot and root tissues. Values are means ± SD (n=3). Asterisks indicate statistically significant difference between wild type and the *zmccd8* mutant (P<0.01, t-test). (C) Venn diagrams showing the comparisons made between the two RNA-seq datasets used in this study. (D) Differentially expressed genes in root tissues overlapping the two datasets. (E) Expression levels of *IRO2;1* in wild-type and mutant siblings growing in iron-sufficient or iron-deficient conditions. Significance was determined using Student’s *t*–test (unpaired, unequal variance). Asterisks (*) indicate significance (* p < 0.05, ** p < 0.01, *** p < 0.001, **** p < 0.0001).

Using our previously generated iron-regulated gene list, we identified all genes that both respond to iron deficiency and are mis-regulated in *ccd8*. In the roots, thirty iron-regulated genes were mis-regulated in *ccd8*, while in the shoots there was only one (Figure 5C). Among these 30 genes is *IRO2;1*, which encodes a predicted transcriptional regulator that is important in activating expression of iron uptake. The observation that ∼3% of the genes that are mis-regulated in *ccd8* mutants are iron-regulated genes (30/1,066), while only about 0.7% of the genes in the total genome (216/32,000) are iron-regulated suggests that iron-regulated genes are over-represented among the *ccd8* mis-regulated genes in roots (hypergeometric p-value = 4.23^11^). Twenty-two of the thirty genes that were down-regulated in the *ccd8* mutant roots are normally upregulated in roots during iron deficiency (Figure 4D and Table 1). Thus, in the strigolactone-less mutant *ccd8*, genes associated with iron uptake are expressed at abnormally low levels. In the shoots, the one gene identified as both iron-regulated and SL-regulated was *ZmYS1*. *ZmYS1* is upregulated in the shoots during iron deficiency, but in the *ccd8* mutant shoots *ZmYS1* was strongly suppressed (FC=- 4.62). Overall, for both roots and shoots, expression of these genes in *ccd8* mutants is opposite to the pattern of expression that occurs during iron deficiency.

**Figure 5:**
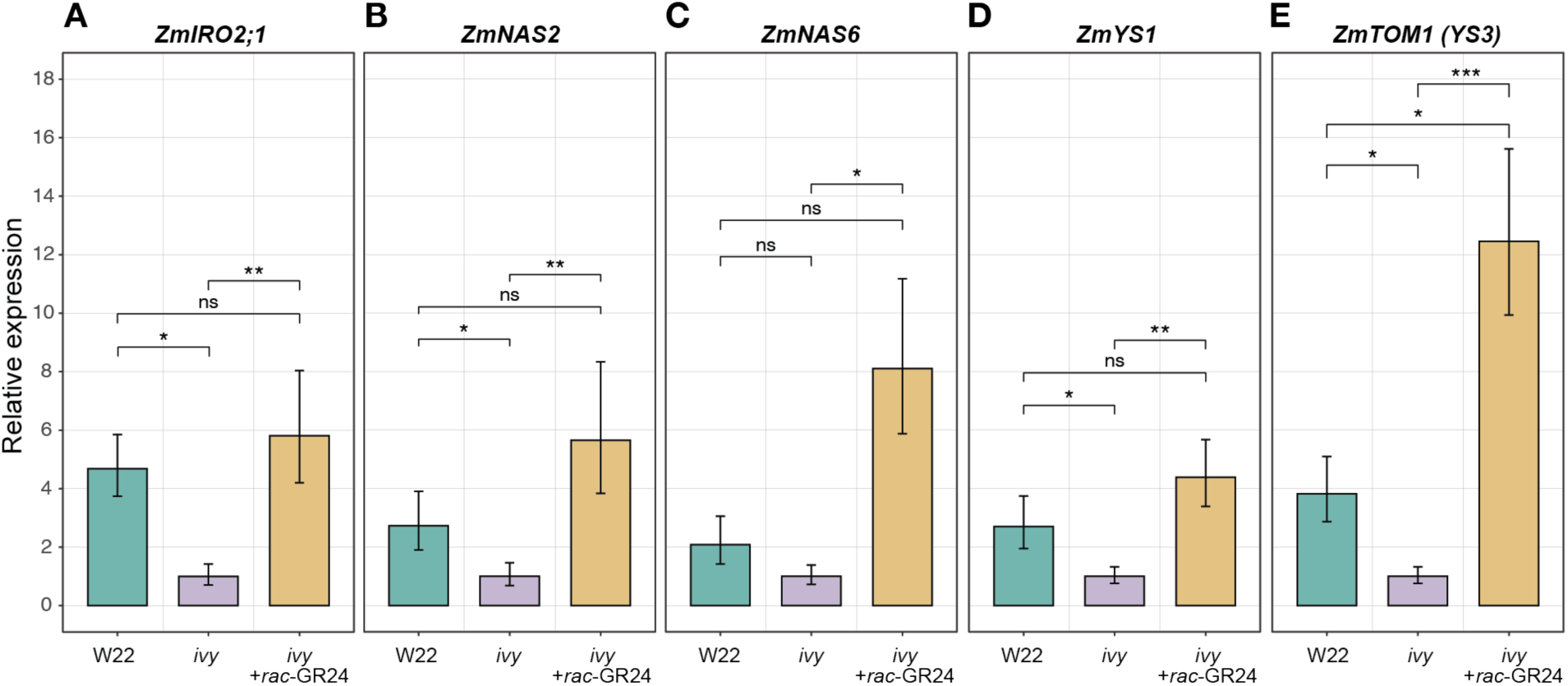
The effect of external SL application on iron-regulated gene expression in *ivy* mutant plants. W22 and *ivy* plants were grown in hydroponic culture. One set of *ivy* plants was supplemented with *rac*-GR24 in the hydroponic solution (*ivy* + rac-GR24), while the other set of *ivy* plants received mock treatment (*ivy*). Gene expression was measured using RT-qPCR. Statistical analysis was performed using one-way ANOVA with posthoc Tukey HSD. Asterisks (*) indicate significance (* p < 0.05, ** p < 0.01, *** p < 0.001, **** p < 0.0001).

**Table 1.**

| Gene ID | Gene product | Fold change<br>[+Fe vs -Fe] | Fold change<br>[WT vs *ccd8*] |
| --- | --- | --- | --- |
| Zm00001eb140680 | IRO2;1 – iron-related transcription factor 2;1 | 6.92 | -3.11 |
| Zm00001eb024180 | haloacid dehalogenase-like hydrolase domain-containing protein | 4.44 | -2.05 |
| Zm00001eb193940 | HSP22 – heat shock protein22 | 4.06 | -2.50 |
| Zm00001eb314230 | GEX1 – gamete expressed1 | 3.97 | -2.40 |
| Zm00001eb074080 | prolyl endopeptidase family protein | 3.48 | -3.00 |
| Zm00001eb341000 | HLH DNA-binding domain-containing protein | 3.31 | -2.08 |
| Zm00001eb077640 | threonine aldolase family protein | 2.83 | -2.20 |
| Zm00001eb333000 | PER24 – peroxidase24 | 2.79 | -2.69 |
| Zm00001eb322710 | hypothetical protein, unknown function | 2.74 | -2.44 |
| GRMZM2G035599 | RPI1 – ribose 5-phosphate isomerase1 | 2.69 | -2.04 |
| Zm00001eb340550 | THX47 – trihelix-transcription factor 47 | 2.59 | 2.24 |
| Zm00001eb014700 | NAS2;1 – nicotianamine synthase2;1 | 2.53 | -3.64 |
| Zm00001eb014680 | NAS2;2 – nicotianamine synthase2;2 | 2.50 | -3.13 |
| Zm00001eb069350 | Diacylglycerol kinase (DAG kinase) | 2.48 | 3.39 |
| Zm00001eb161140 | membrane protein, unknown function | 2.43 | -5.79 |
| Zm00001eb396120 | NAS6;2 – nicotianamine synthase6;2 | 2.31 | -5.31 |
| Zm00001eb010040 | DMAS1 – deoxymugineic acid synthase1 | 2.28 | -4.57 |
| Zm00001eb113920 | TAT1 – tyrosine aminotransferase1/ NAAT-L1<br>nicotianamine aminotransferase-like1 | 2.23 | -3.57 |
| Zm00001eb396110 | NAS6;1 – nicotianamine synthase6;1 | 2.20 | -4.42 |
| Zm00001eb231210 | NRAMP6/ NRAT2 – nramp metal transporter/<br>nramp aluminum transporter2 | 2.18 | -2.19 |
| Zm00001eb058070 | CYP33 – cytochrome P450 CYP81A39 | 2.15 | 2.00 |
| Zm00001eb383440 | E3 ubiquitin-protein ligase | 2.15 | -4.22 |
| Zm00001eb195880 | inositol polyphosphate phosphatase (Synapto-<br>genin-like) family protein | 2.09 | -7.10 |
| Zm00001eb410380 | RBOH7 – respiratory burst oxidase7 | 2.08 | 2.25 |
| Zm00001eb203620 | xyloglucan endotransglucosylase/hydrolase pro-<br>tein26 | 2.06 | -3.27 |
| Zm00001eb193370 | PP67 – tyrosine-protein phosphatase67 | 2.02 | -2.23 |
| Zm00001eb249940 | Aquaporin PIP1-2 | -2.04 | -5.68 |
| Zm00001eb235330 | DUF642 – domain of unknown function 642<br>containing protein | -2.17 | -4.21 |
| Zm00001eb419530 | EXPA3 – alpha expansin3 | -2.31 | -3.50 |
| Zm00001eb111420 | PER66 – peroxidase66 | -4.41 | -2.40 |

Next, we asked whether *ivy* mutants appropriately alter gene expression in response to limited iron supply in the growth medium. Plants from an *ivy-*segregating population were grown in hydroponic culture, and iron was withheld from half the plants. In both *ivy* mutant and WT roots, growth in -Fe medium caused a similar marked increase in the level of *ZmIRO2;1* mRNA (Figure 4E). Likewise, *ZmIRO2;1* mRNA increased after growth in -Fe medium similarly in both *ivy* mutants and WT shoots. This result indicated that *ivy* mutants are able to respond vigorously to exogenously imposed iron-deficiency, *i.e.*, withdrawal of iron from the growth medium, even though basal levels of iron-responsive genes are abnormally low in *ivy* plants during growth in normal, iron sufficient medium or soil.

### SL application causes altered expression of iron-regulated genes

We tested whether the external provision of synthetic SL to *ivy* mutant plants could increase expression of iron-regulated genes to wild-type levels. We grew *ivy* and wild-type W22 plants in hydroponic culture. One set of *ivy* plants was provided with *rac*- GR24 in the hydroponic media, while the control *ivy* plants and the W22 wild-type control plants did not receive *rac*-GR24. We then measured the expression of *ZmIRO2;1*, *ZmNAS2*, *ZmNAS6*, *ZmYS1* and *ZmTOM1* in the roots. Although RNA-seq analysis did not indicate that *ZmYS1* or *ZmTOM1* are significantly down-regulated in *ccd8* roots, we included them here because they are likely downstream targets of ZmIRO2;1. As expected from the RNA-seq results, *ZmIRO2;1* expression in *ivy* roots was more than 4-fold lower than wild-type. Notably, *ivy* plants that were treated with *rac*- GR24 expressed *ZmIRO2;1* at levels indistinguishable from wild-type (Figure 5). The results were similar for *ZmNAS2, ZmNAS6 and ZmYS1*. For *ZmTOM1*, the expression became significantly higher than wild-type in the presence of *rac*-GR24. These findings support the idea that SLs affect the expression of these iron-regulated genes.

### SL mutants in rice and Arabidopsis

The idea that mutation of *ccd8* causes iron deficiency only in maize was puzzling, but although *ccd8* mutants have been characterized in both rice and Arabidopsis, chlorosis was not noted in these mutants (Auldridge et al., 2006; Arite et al., 2007). We re-examined *ccd8/d10* and *d14* mutants of rice and *ccd8/max4* mutants of Arabidopsis. When the rice mutants were grown in normal culture, we were able to observe an obvious chlorotic phenotype relative to WT control plants (Figure 6A). The Arabidopsis mutants did not exhibit visible chlorosis when grown in soil, but when they were grown on plates with 10 uM iron instead of the normal 100 uM iron, they showed pronounced chlorosis (Figure 6B, C and D). WT control plants grown in the same 10 uM iron condition were not chlorotic (Fig 6B and C.) This indicates that the effect of SL hormone on iron homeostasis is not unique to maize, but also occurs in other species.

**Figure 6.**
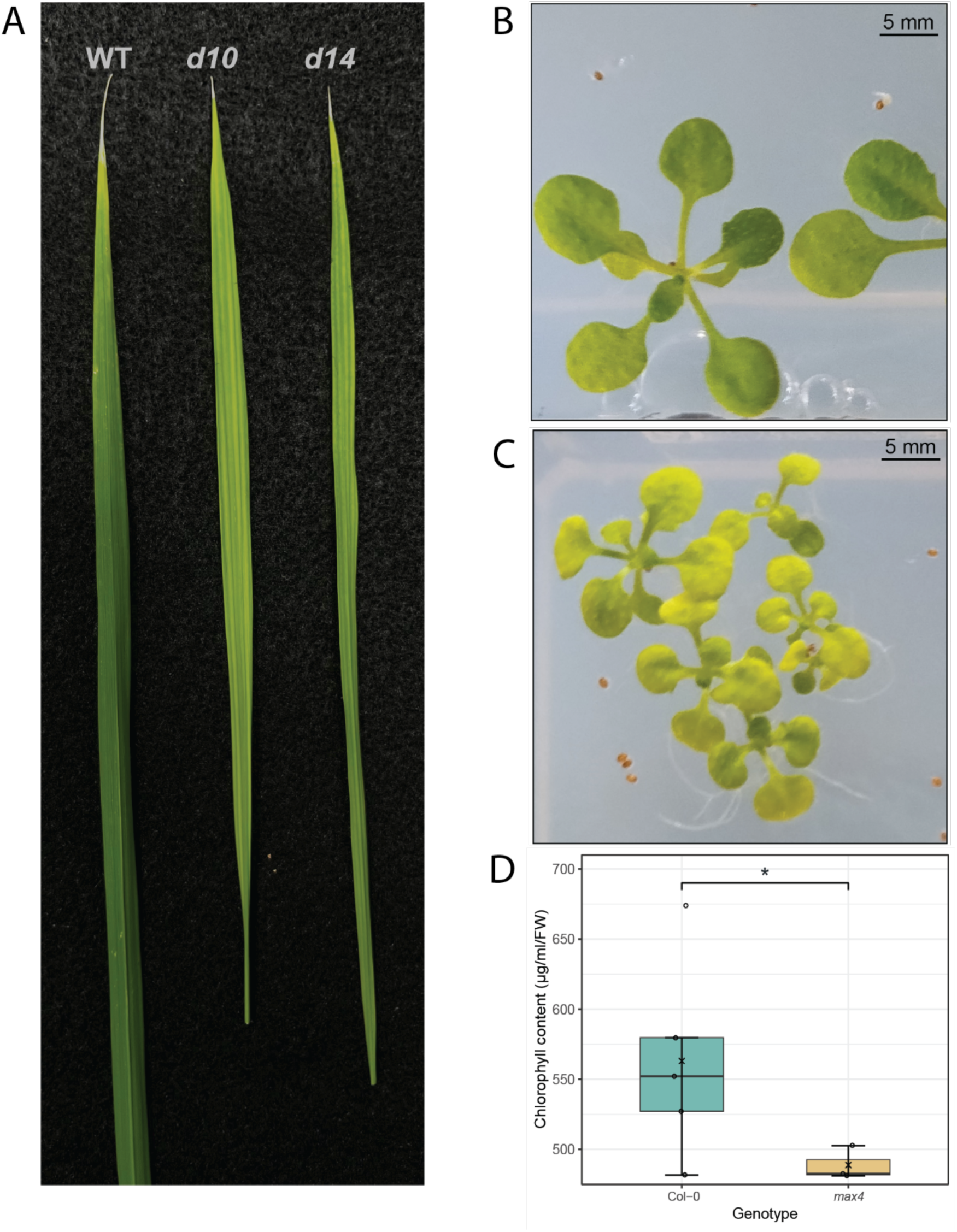
Phenotypes of strigolactone mutants in rice and Arabidopsis. (A) Leaves of rice *d10* (*ccd8*) and *d14* mutants compared to wild-type (background “Shiokari”). (B) Arabidopsis Col-0 growing in 10 μM iron. (C) Arabidopsis max4 (*ccd8*) mutants growing in 10 μM iron. (D) Chlorophyll concentration of shoots from Arabidopsis plants grown in 10 μM iron. Significance was determined using pairwise Student’s *t*-test (unpaired, unequal variance). Asterisks (*) indicate significant difference between wild-type and mutant (* p < 0.05, ** p < 0.01, *** p < 0.001, **** p < 0.0001).

## Discussion

In this study, we characterized a novel iron-deficient maize mutant, *interveinal yellow* (*ivy*). The *ivy* mutant responds particularly well to foliar iron application (Chan-Rodriguez and Walker, 2018), thus making it an attractive candidate for identifying a new gene involved in iron homeostasis. Using BSA-seq, we showed that the mutated locus responsible for the iron-deficient phenotype encodes CCD8, a key enzyme in SL biosynthesis (Figure 1). Complementation crosses with a previously characterized *ccd8* mutant (Guan et al., 2012) confirmed this finding (Figure 2). The identification of an iron-deficient mutant as an SL mutant indicated that there is a possible connection between this class of hormones and iron homeostasis in maize. The yellow-striped phenotype of a maize *ccd7* mutant identified in this study further confirms that genetically impaired biosynthesis of SL leads to iron deficiency symptoms in maize. Maize is not the only species to show this phenomenon. In SL mutants of both rice and Arabidopsis, chlorosis is apparent in mutants growing in conditions where the WT does not show chlorosis.

SLs have well established roles in coordinating plant growth and development in response to nitrate and phosphate conditions. SL exudation from roots is stimulated by phosphate and nitrate deficiency as a signal that promotes formation of arbuscular mycorhizzae (Yoneyama et al., 2007b; Yoneyama et al., 2007a). In addition, changes in root architecture that occur in response to nutrient availability depend on SL activity (Arite et al., 2012; Sun et al., 2014; Sun et al., 2021). Levels of intermediary metabolites also affect SL levels or SL signaling in plants (Bertheloot et al., 2020; Tal et al., 2022). All these phenomena indicate tie-ins between SLs and the maintenance of appropriate plant growth according to abiotic conditions (recently reviewed in (Barbier et al., 2023). But in these phenomena, the direction of signaling is from nutrient status to an alteration in SL hormone levels. Here, we describe tie in between a plant nutrient and SL hormones that operates in the opposite direction. SL hormones exert control on the uptake of the essential nutrient iron.

### Evaluation of the maize *ccd8* mutant phenotype

To begin to understand how a mutant CCD8 enzyme can cause an iron-deficient phenotype in maize, we considered several distinct hypotheses. First, we posited that CCD8 could have a second, poorly understood enzymatic activity outside of the SL biosynthetic pathway. The observation that supplying synthetic SL to mutants alleviated chlorosis indicated that the iron-deficient phenotype is caused specifically by the lack of SLs (Figure 3). The chlorotic phenotype of a second SL biosynthetic mutant, *ccd7*, also corroborates this idea.

Next, we hypothesized that altered root crown root development in *ccd8* mutants (Guan et al., 2012) led to impaired iron uptake, even though the iron deficient phenotype displays at an early seedling stage before crown root development. It has been suggested that this crown roots might have a particular role in nutrient uptake (Blizard and Sparks, 2020), and further, we noticed that expression of key iron-regulated genes is strongest in the crown roots (Stelpflug et al., 2016) However other maize mutants that are impaired in crown root development *rt1* and *rtcs,* do not exhibit chlorosis in above ground parts, indicating that the absence of crown roots does not explain the iron deficiency chlorosis of *ccd8* mutants.

Instead, our data indicate that SL modulates the expression of iron deficiency associated genes, particularly those that have increased expression during iron deficiency. The expression of such genes is low in *ccd8* mutants (Fig 4). Application of the synthetic SL hormone rac-GR24, increases expression of these genes (Fig 5), and, importantly, application of rac-GR24 to ccd8 mutants leads to greening of the ordinarily chlorotic mutant leaves (Fig 3). Thus, we conclude that SLs modulate the expression of gene involved in iron uptake.

### Transcriptomic evaluation of iron-deficient conditions in maize

As part of this study, we evaluated the transcriptomic changes that take place in maize roots and shoots under sufficient, deficient and resupplied iron conditions. Through this analysis, we established a set of iron-regulated genes for roots and shoots in maize, and categorized each iron-regulated gene as “fast-responding” “slow-responding” after a period of 24 hours of iron resupply. The fast-responding subset included genes with known roles in iron homeostasis, such as *IRO2;1*, *YS1*, *TOM1* and *NAS* genes. Only seven genes were iron-regulated in both roots and shoots including *IRO2;1* and *YS1*. Gene ontology enrichment analysis of differentially expressed genes in the roots revealed their involvement in processes such as NA and PS biosynthesis, and iron transport and homeostasis (Supplemental data 3). This implies that the primary response of roots to limited iron is the promotion of iron uptake mechanisms. In shoots, the main processes affected in were photosynthetic and porphyrin-related metabolic processes, and these genes were all down-regulated (Supplemental data 3). This suggests that the primary response of shoots to limited iron is the suppression of photosynthetic and metabolic reactions.

Similar RNA-seq experiments performed in Arabidopsis identified numbers of root iron-responsive genes similar to that in maize, with 191 genes increasing in abundance and 40 genes decreasing in abundance (Grillet et al., 2018). In Arabidopsis shoots, the number of genes that increased in abundance was similar to that in maize, but the number of genes with decreased expression during iron deficiency was almost 3 times that in maize (Bakirbas and Walker, 2022). In both Arabidopsis and rice, iron homeostasis is regulated by several members of the basic helix-loop-helix (bHLH) family of transcription factors (reviewed recently by (Liang, 2022; Gao et al., 2024)). A key regulator of iron uptake is FIT, first identified in tomato as FER (Ling et al., 2002; Colangelo and Guerinot, 2004; Liang et al., 2020)). *FIT* expression is induced by iron deficiency in roots and FIT upregulates iron uptake-related genes, including *IRT1* and *FRO2* in Arabidopsis or *OsYSL15* and PS biosynthesis-related genes in rice (Liang et al., 2020; Wang et al., 2020b) in rice. We observed two *FIT* homologs in maize, *ZmbHLH100* (*FIT1;1 Zm00001eb420910*) and *ZmbHLH101* (*FIT1;2 Zm00001eb085690*). *FIT1;2* is upregulated 2.6-fold in roots during iron deficiency, and was also noted as being down-regulated in *ccd8* mutants, although, at the FDR <0.01 used in our analysis, this was not significant. *FIT1;1* is more moderately upregulated (1.5-fold) during iron deficiency, and was not detected during RNAseq analysis of *ccd8* mutants. FIT forms heterodimers with subgroup Ib bHLHs (bHLH38/39/100/101 in Arabidopsis; IRO2 in rice), which are also iron-regulated (Wang et al., 2007; Yuan et al., 2008; Grillet et al., 2018). (Carretero-Paulet et al., 2010; Liang et al., 2020; Wang et al., 2020b). As shown here, *OsIRO2* has two orthologs in maize, *ZmIRO2;1* and *ZmIRO2;2*, with *ZmIRO2;1* showing a strong response under iron-deficient conditions, and strong down-regulation in *ccd8* mutants.

### SL involvement in iron homeostasis is not specific to maize

Mutants that are incapable of producing SL have been studied in both rice and Arabidopsis, but in those species chlorotic phenotypes were not reported. We re-examined the rice *d10* (*ccd8*) and *d14* (*SL receptor*) mutants, and observed that both exhibit chlorosis in normal culture conditions. This is very similar to the observations we have made regarding the *ccd8* and *ccd7* mutants in maize, and suggests that in rice, too, the inability to make or sense SL hormones leads to iron deficiency. In Arabidopsis, we did not observe chlorosis when plants were grown on soil or when they were grown on MS plates containing 100 uM iron. However, when the plants were grown on plates with a reduced level of iron (10 uM), *ccd8* (*max4*) mutants were small and showed pronounced chlorosis. WT plants grown in the same conditions were green and had higher chlorophyll. This indicates that the connection between SL hormone and iron is likely to exist in many plant species including both monocots and dicots.

In maize, the product of CCD8 enzymatic activity, carlactone, in converted either to carlactonic acid (CLA) by ZmMAX1b or to 3-oxo-19-hydroxy-carlactone by ZmCYP706C37. By changing the flux into the 3-oxo-19-hydroxy-carlactone side of the pathway using mutation of *ZmMax1b*, the SL profile can be shifted towards production of zealactol and zealactonic acid and away from production of zealactone. This has a pronounced effect on germination of Striga, which is stimulated by zealactone (Li et al., 2023). Maize *max1b* mutants do not show iron deficiency chlorosis, suggesting that this shift in SL profile away from zealactone does not have deleterious affects on iron homeostasis. The identity of the SL(s) that are needed for proper iron homeostasis are not currently clear.

In Arabidopsis, gene expression after treatment with the SL analog rac-GR24 has been examined (Wang et al., 2020a). Further, the sites of binding of a key negative regulator involved in SL signaling, SMXL6, have been examined using ChIP-seq (Wang et al., 2020a). Interestingly, the Arabidopsis *FIT1* gene, which is well characterized as an essential component in iron deficiency gene regulation, is not only down-regulated following rac-GR24 treatment, but also, SMXL6 binds to the upstream portion of the *AtFIT1* (Wang et al., 2020a). These observations strongly suggest a model for SL regulation of iron homeostasis, as illustrated in Suppl. Fig. S7. In the absence of SL, no complex is formed between the SL receptor D14, the ubiquitin ligase SCF^max2^, and the SMXL6 negative transcription regulator. This allows SMXL6 to interact with the *AtFIT1* gene, suppressing its transcription, which in turn is expected to cause lower basal iron uptake. In *ccd8* mutants, the lack of SL causes permanently lowered iron uptake, leading to chlorosis. In the presence of SL hormones, the complex hormone receptor complex is formed, SMXL6 is degraded, and *FIT1* is released from repression.

Our transcriptomic data on the maize *ccd8* mutant indicated that the gene encoding the presumed binding partner of *ZmFITs*, *ZmIRO2;1*, is the most clearly down-regulated transcription factor gene in *ccd8* mutants. Neither of the two maize *FIT* genes had significantly decreased (log_2_(|FC|) ≥ 1;FDR adjusted p-value ≤ 0.01) expression in *ccd8* plants, nor did the second *IRO2* gene of maize (*ZmIRO2;2)* have significantly decreased expression in *ccd8* plants. We hypothesize a model in which the target of SL regulation in maize is not *FIT1*, as in Arabidopsis, but instead is its binding partner, *IRO2* (Suppl Fig S6).

### Iron deficiency and basal iron uptake

The iron deficient, chlorotic phenotype of maize *ccd8* mutants presents a conundrum. Why do *ccd8* plants not sense ‘iron deficiency’ during growth in normal iron conditions? We have shown that they are capable of mounting a normal transcriptional response to externally imposed iron deficiency, after all. The mechanism through which plants sense iron deficiency remains mysterious. The BTS and BTSL (HRZ in rice) proteins are candidates for this sensing function (Long et al., 2010), but current models indicate that their role is to limit Fe over-accumulation (Rodríguez-Celma et al., 2019) and to shut down iron accumulation during bacterial infection (Cao et al., 2024). Additional ‘sensors’ may exist to initiate iron uptake during deficiency. Further, the location(s) of iron deficiency sensing are not clear. Local transcriptional control of iron uptake has been extensively documented (Liang, 2022), but long distance, shoot-based sensing clearly also occurs (Mendoza-Cozatl et al., 2014; Zhai et al., 2014; Gayomba et al., 2015). In the absence of more information about the mechanisms(s) of iron sensing, it is difficult to understand the reasons why *ccd8* plants do not sense their own internal iron deficiency and adjust gene expression to correct it.

## Materials and methods

### Plant material and growth conditions

The *ivy* mutant seed stock was obtained from the Maize Genetics Cooperation Stock Center (MGCSC #3812O; *ys*^∗^-*N2398*). The *ccd8::Ds* mutant allele was originally developed in a W22 inbred (Vollbrecht et al., 2010; Guan et al., 2012) and was subsequently introgressed into B73 inbred background by 6 generations of back-crossing. The mutant stock for *ccd7* was obtained from the Mu-Illumina project (mu-illumina_578516.6) (Williams-Carrier et al., 2010). The *rt1* and *rtcs1-1* mutant stocks were obtained from MGCSC (#311E and #131H, respectively). The *ys1:ref* and *ys3:ref* stocks were obtained from MGCSC (#503A and #311F, respectively) and have been maintained in homozygosity for several generations. The Arabidopsis *ysl1ysl3* double mutant was previously generated from the Salk T-DNA insertion mutants *ysl1-2* (SALK_034534) and *ysl3-1* (SALK_064683) and maintained in homozygosity (Waters et al., 2006). The *max4-6* mutant stock was previously described (Auldridge et al., 2006) and maintained in homozygosity. The wild-type *Arabidopsis thaliana* accession Columbia-0 (Col-0) was used as control.

For growth in the soil, maize or Arabidopsis seeds were planted in plastic pots with a 4:1 mixture of soil (Sun Gro^®^ Professional Growing Mix) and Turface® MVP®, pretreated with Gnatrol (Valent BioSciences). Rice plants were planted I plastic pots with an equal mixture of soil and Profile® Greens Grade™. Arabidopsis plants were grown at room temperature (20–25 °C) with 16 hours of light and 8 hours of dark. Maize and rice seeds were grown in the greenhouse with 16 hours of light and 8 hours of dark. For growth on plates, Arabidopsis seeds were sterilized (70% ethanol, 0.05% Triton X- 100) and grown on sterile culture plates as described previously (Bakirbas and Walker, 2022). For growth in hydroponics, maize seedlings were grown as described previously (Fili and Walker, 2023). Throughout this article, the term “hydroponic culture” refers to plants growing in solution with normal iron concentration and the term “iron-deficient hydroponic culture” refers to plants growing in solution with no added iron. For supplementation of synthetic SL in the hydroponic culture (+/-)-GR24 (*rac*-GR24) was used (PhytoTech Labs, G3324) at a 1 μM final concentration (Guan and Koch, 2020). The hydroponic solution was renewed every 3 days.

### Morphological measurements

Morphological measurements in *ivy* mutants were performed in siblings from F2 populations generated by crossing *ivy* to W22 or B73. The W22 population consisted of 107 individuals (25 mutant, 82 wild-type) and the B73 population consisted of 139 individuals (39 mutant, 100 wild-type). Plants were grown in the field and all measurements were performed when the plants reached maturity.

For quantification of crown root emergence in the *ivy* mutants, seedlings were grown in hydroponic culture. One set of *ivy* plants was supplemented with *rac*-GR24 every 3 days in the hydroponic solution (*ivy* + rac-GR24), while the other set of *ivy* plants received mock treatment (*ivy*). The length of each crown root was measured on day 9. Six biological replicates were used per genotype.

### DNA and RNA isolation

To obtain DNA for genotyping of *ivy* mutants and Sanger sequencing a modified version of the Dellaporta miniprep was used (Dellaporta et al., 1983). For genotyping of *ccd7* mutants using TaqMan assays, a modified quick DNA extraction protocol was used (Wang et al., 1993).

To obtain RNA for gene expression measurements with qRT-PCR, root or shoot tissues were ground in 2 mL tubes (Qiagen) containing 3 chrome steel beads of 3.2 mm diameter (BioSpec Products) and using either the TissueLyser II (Qiagen) or the Mixer Mill MM 400 (Retsch) with pre-frozen blocks. Total RNA was extracted using the RNeasy Plant Mini Kit (Qiagen) with on-column DNAse treatment. At least 3 biological replicates were used per genotype or condition for all gene expression measurements. RNA concentration was quantified using a NanoDrop (ThermoScientific).

### cDNA synthesis and quantitative real-time PCR

For gene expression measurements in *ivy* mutants, cDNA was synthesized from at least 500 ng of total RNA using SuperScript IV VILO Master Mix (Invitrogen™, Thermo Fisher Scientific). qRT-PCR reactions were set up using the PowerUp™ SYBR™ Green Master Mix (Applied Biosystems™, Thermo Fisher Scientific) in MicroAmp™ Fast Optical 96-Well Reaction Plates (Applied Biosystems™, Thermo Fisher Scientific). The reactions were run on a StepOnePlus™ Real-Time PCR System (Applied Biosystems™, Thermo Fisher Scientific). Relative expression was determined using the ΔΔC_T_ method with C_T_ values of target genes normalized to *ZmGAPDH*.

For additional analysis of key transcripts in RNA-seq profiles of *ccd8*, 5.5 μg of the same total RNA used for the RNA-seq libraries were first treated with RQ1 RNase-free DNase (Promega) to confirm (via negative controls) that there was no carryover of genomic DNA. For quantitative PCR, a Power SYBR green RNA-to-C_T_, 1-step kit (Applied Biosystems) was used with an iCycler iQ real-time PCR detection system (Bio-Rad). The *18S rRNA* was used an internal reference. Primer sequences were adapted from (Nozoye et al., 2013). Statistical significance of differential expression was tested by ANOVA.

### BSA-seq analysis

The original *ivy* stock was crossed to the W22 inbred. After self-crossing the F1 generation, a segregating F2 population was generated. The population was grown in the field and scored for the yellow stripe phenotype. Leaf collection was performed from a total of 120 yellow stripe individuals and 120 wild-type individuals. Four punches were obtained from each leaf and pooled together for a bulk DNA extraction. To obtain high-quality genomic DNA for BSA-seq, the CTAB method for DNA extraction was used as described previously (Healey et al., 2014). The extracted DNA was quantified using Nanodrop and Qubit and quality was assessed using gel electrophoresis. Library preparation and sequencing were performed by Novogene (Hi-seq 4000, Illumina). The quality of the raw reads was assessed using FastQC (Andrews, 2010b). Adapter sequences and low-quality reads were removed using Trimmomatic (Bolger et al., 2014) with an average quality cutoff of 24. Trimmed reads were aligned to the maize W22v2 genome (assembly version: *Zm-W22-REFERENCE-NRGENE-2.0*) using the Bowtie2 aligner (Langmead and Salzberg, 2012). Variant calling was performed using SAMtools mpileup and variants were filtered using a base quality above 20 and a base coverage above 8 (Li et al., 2009). Allele frequency was calculated and plotted using custom R scripts. Variants were also identified using Varscan (Koboldt et al., 2012) for downstream SNP analysis. All SNPs within the mapping region were extracted and filtered only for homozygous in the mutant pool. SNPs that were also homozygous in the wild-type pool were filtered out. Naturally occurring variants were further filtered out using the available genomes of the following maize inbreds: A632, B73, B104, EP1, F7, Mo17, Oh43 and PH207 (Hirsch et al., 2016; Jiao et al., 2017; Springer et al., 2018; Sun et al., 2018; Haberer et al., 2020; Wang et al., 2023). The region corresponding to the mapping interval in W22 was identified in each inbred through synteny and multiple alignment was performed using Mauve (Darling et al., 2004) and setting the W22 region as reference. The polymorphisms of all the inbreds were extracted and if they matched the homozygous variants identified in the *ivy* pool they were removed. The effect of each of the remaining SNPs on the protein product was analyzed using SNPeff (Cingolani et al., 2012), identifying moderate and high-effect SNPs. For moderate-effect SNPs their impact on the protein function was further analyzed using PROVEAN (Choi and Chan, 2015). All genome assembly and annotation files were retrieved directly from MaizeGDB (maizegdb.org).

### RNA-seq analysis

For transcriptomic analysis of iron deficiency and resupply, wild-type plants (W22) were grown in hydroponic culture for 12 days and then switched to +Fe or −Fe for 3 days to induce an iron deficiency environment. After the 3 days, one set of −Fe plants was resupplied with hydroponic solution containing 0.1mM FeSO_4_-EDTA (±Fe plants) while the +Fe and −Fe plants received new solution. After 24 hours, the roots and shoots were sampled separately and immediately frozen in liquid nitrogen. Three biological replicates were used for each condition. The tissues were ground using pre-chilled mortars and pestles. The total RNA from roots and shoots was isolated using the Direct-zolTM RNA MiniPrep Kit (Zymo Research) with DNase I treatment according to the manufacturer’s instructions. The extracted RNA was quantified and assessed for quality using an Agilent 2100 Bioanalyzer. Library preparation and sequencing were performed by Novogene (Hi-seq 4000, Illumina). The adapter sequences and low-quality reads were removed using Trim Galore (Andrews, 2010a; Martin, 2011; Krueger et al., 2023). Trimmed reads were aligned to the maize W22v2 genome (assembly version: *Zm-W22-REFERENCE-NRGENE-2.0*) using the HISAT2 aligner (Kim et al., 2015). Transcript assembly and quantification was performed using StringTie (Pertea et al., 2015). Differential gene expression analysis was performed using DESeq2 (Love et al., 2014). The cutoff criteria that were used were log_2_(|FC|) ≥ 1 and Benjamini-Hochberg (FDR) adjusted p-value ≤ 0.01 (Benjamini and Hochberg, 1995). Genome assembly and annotation files of the W22v2 genome were retrieved directly from MaizeGDB (maizegdb.org).

For analysis of expression profiles of *ccd8* mutants, both wild-type B73 and *ccd8* mutant seedlings were grown in plastic containers (diameter=11.2 cm & height=9.4 cm) with propagation mix for growth media (Sun Gro^®^). The growth chamber environment was set at 25°C with a diurnal cycle of 14-hour light (150 µmole m^-2^ s^-1^) and 10-hour dark. Root and shoot tissues were sampled from seedlings at 14 days after planting immediately frozen in liquid nitrogen, and stored at −80°C until use. Tissues were ground in liquid nitrogen prior to extraction and purification of total RNA by plant RNeasy (Qiagen). The resulting RNA was quantified by NanoDrop 1000 (Thermo Fisher Scientific). Libraries for RNA-seq were prepared with a TruSeq sample preparation kit (Illumina) and sequenced by Hi-seq 2000 (ICBR, University of Florida). The data were analyzed using the Tuxedo Suite with statistical significance of differentially expressed genes (DEGs) set at FDR≤0.01 (geometric normalization). The RNA-seq data for *ccd8* and B73 root tissues are available in the NCBI database (BioProject PRJNA757767) under accession numbers of SRR15613590, SRR15613591, SRR15613599, SRR15613593, SRR15613594, and SRR15613595.

### Chlorophyll measurements

For measurements in Arabidopsis, Col-0, *ysl1ysl3* and *max4-6* stocks were grown in ½× MS+Fe plates for 10 days. Then all genotypes were switched to ½× MS+Fe or ½× MS–Fe for 3 days to induce iron deficiency. After the third day, the rosettes were collected, weighed, and placed into glass tubes with screw caps containing 1 mL of N-Dimethylformamide (DMF). The tubes were left in the dark at 4°C overnight. The absorbance of chlorophyll α (647nm) and chlorophyll β (664nm) was measured using a Thermo Scientific Spectronic 200 UV spectrophotometer. Six biological replicates were used per genotype. The total chlorophyll content was calculated as Chl = 17.9*A*_647_ + 8.08*A*_664.5_ (Inskeep and Bloom, 1985).

For maize seedlings grown in hydroponic culture, a SPAD meter was used (SPAD 502 Plus Chlorophyll Meter 2900PDL, Konica Minolta). Sampling was performed as previously described, by averaging all measurements within a leaf (Süß et al., 2019). Six biological replicates were used per genotype.

### Metal measurements

For plants growing in hydroponic culture, leaves of at least 10 individual plants at the V4 stage were collected. For plants growing in soil, leaves of at least 10 individual plants at V3 stage were collected. The tissues were rinsed multiple times with double-distilled water (ddH_2_O). The samples were dried at 65°C for 72 hours. All genotypes studied in each experiment were grown simultaneously and using the same hydroponic culture or soil batch. Metal concentrations were determined by inductively coupled plasma mass spectrometry (ICP-MS) at the Donald Danforth Plant Research Institute.

### Homology identification

Homologous genes in maize were identified by BLASTing protein sequences from Arabidopsis or rice against the B73 genome (Zm-B73-REFERENCE-NAM-5.0). Arabidopsis sequences were retrieved from The Arabidopsis Information Resource (TAIR) database (www.arabidopsis.org); rice sequences were retrieved from Phytozome (phytozome-next.jgi.doe.gov). Synteny analysis was performed using the SynFind tool on CoGe (genomevolution.org).

### Statistical analyses

All statistical analyses were performed using custom R scripts in RStudio. When comparing the means between two groups, either the Student’s *t*–test (unpaired, unequal variance) was used or the Mann–Whitney test. When comparing the means between three groups, Analysis of Variance (ANOVA) was used with post-hoc Tukey HSD. The “rstatix” package was utilized for statistical analysis (Kassambara, 2020).

---

**Supplemental figure S1:**
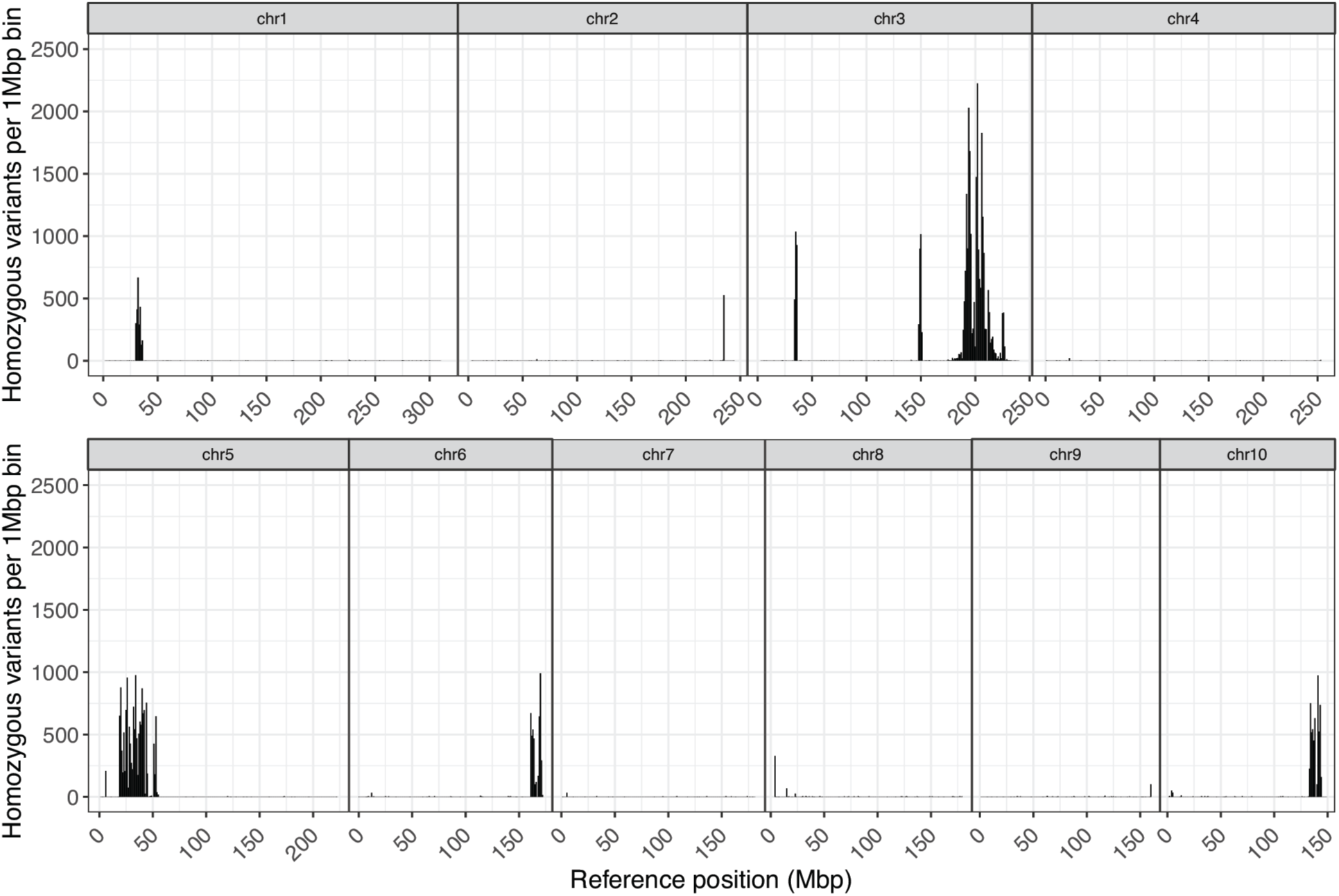
Plots of the number of homozygous variants in the *ivy* mutant pool for chromosomes 1 – 10. Comparison is made against the W22 reference genome (assembly version: *Zm-W22-REFERENCE-NRGENE-2.0*). Homozygous variants that were also present in the wild-type siblings were filtered out. Variants were plotted using 1Mbp bin size.

**Supplemental figure S2:**
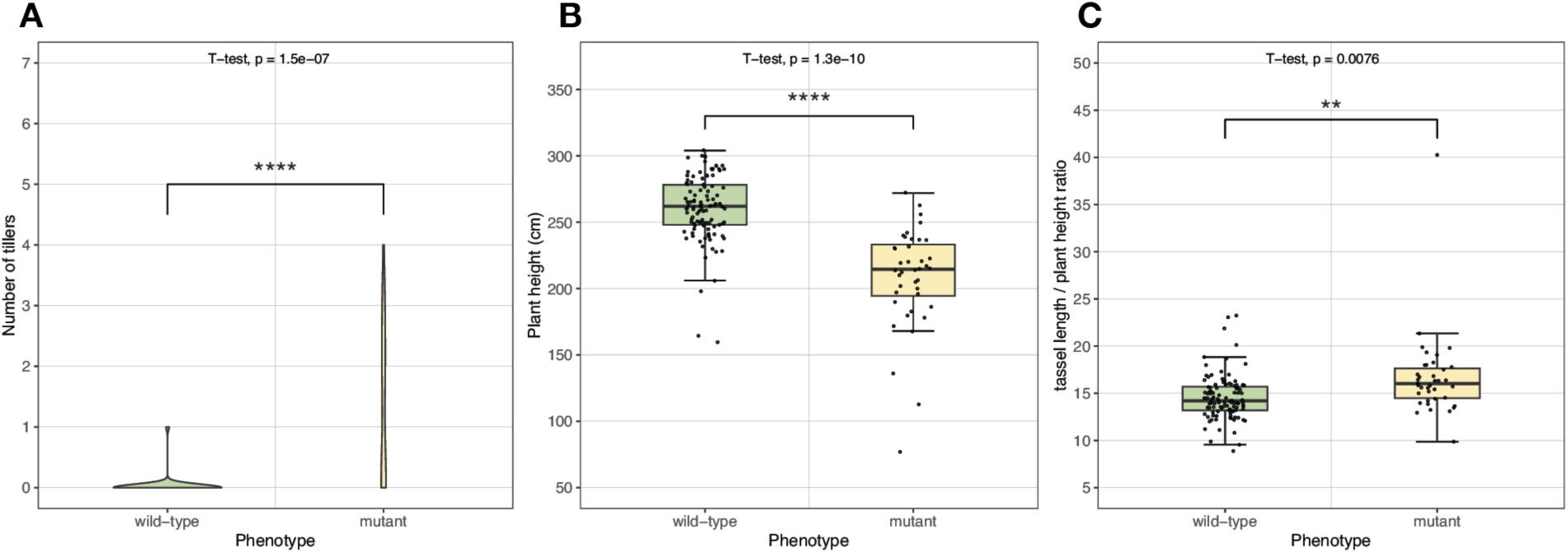
A, B and C, morphological measurements in *ivy* mutants compared to wild-type siblings from an F2 population generated by crossing *ivy* to B73. The population consisted of 139 individuals (39 mutant, 100 wild-type). All measurements were performed when the plants reached maturity. C, Quantification of the total tiller number. D Quantification of the plant height. E, Quantification of the tassel length/plant height ratio. Significance was determined using Student’s *t*–test (unpaired, unequal variance). Asterisks (*) indicate significant difference between wild-type and mutant (* p < 0.05, ** p < 0.01, *** p < 0.001, **** p < 0.0001).

**Supplemental figure S3:**
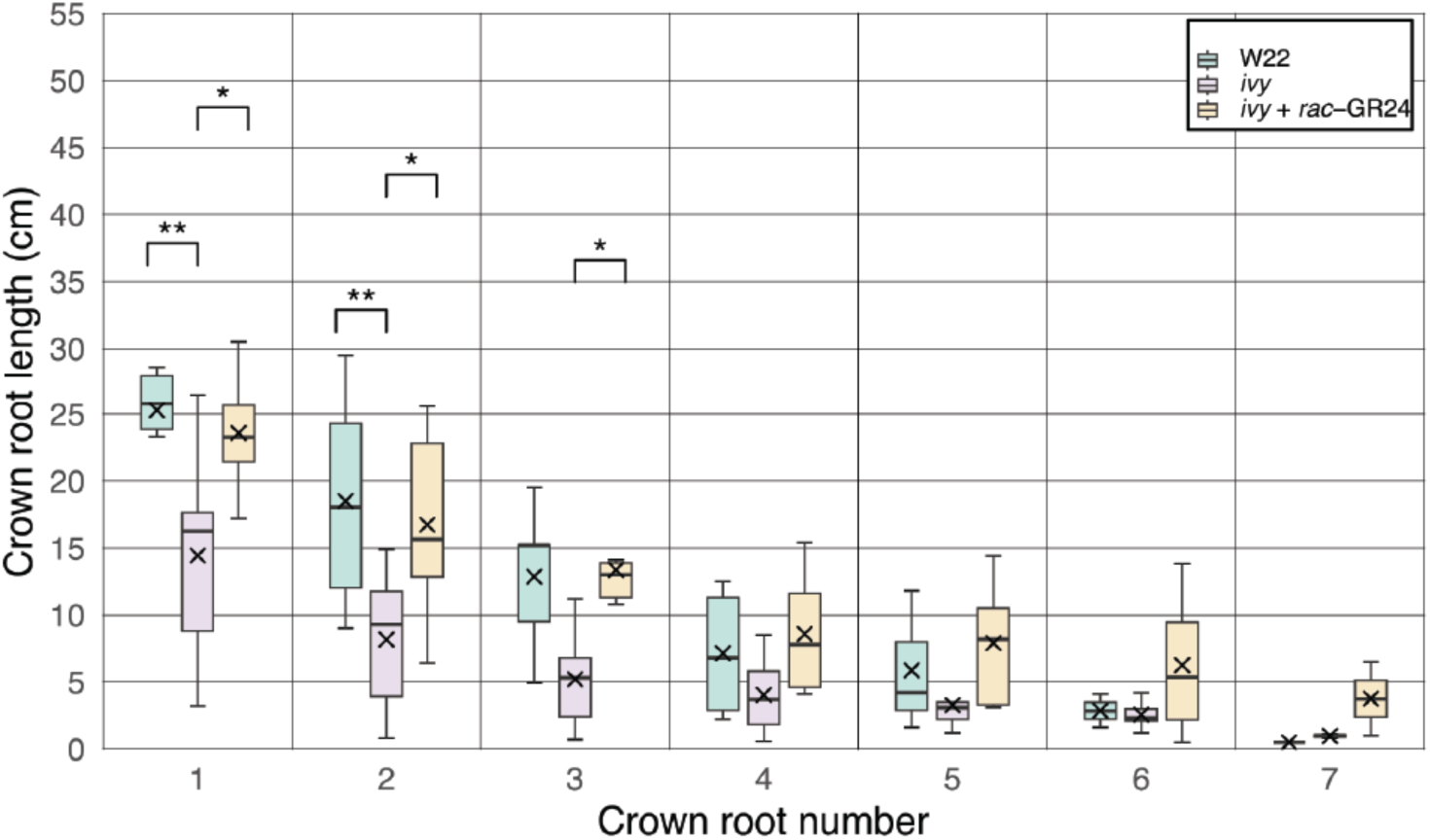
Quantification of crown root emergence in the *ivy* mutant growing in hydroponic culture. One set of *ivy* plants was supplemented with *rac*-GR24 every 3 days in the hydroponic solution (*ivy* + rac-GR24), while the other set of *ivy* plants received mock treatment (*ivy*). The length of each crown root was measured on day 9 (n=6). Statistical analysis was performed using one-way ANOVA with posthoc Tukey HSD. Asterisks (*) indicate significant difference between wild-type and mutant (* p < 0.05, ** p < 0.01, *** p < 0.001, **** p < 0.0001).

**Supplemental figure S4:**
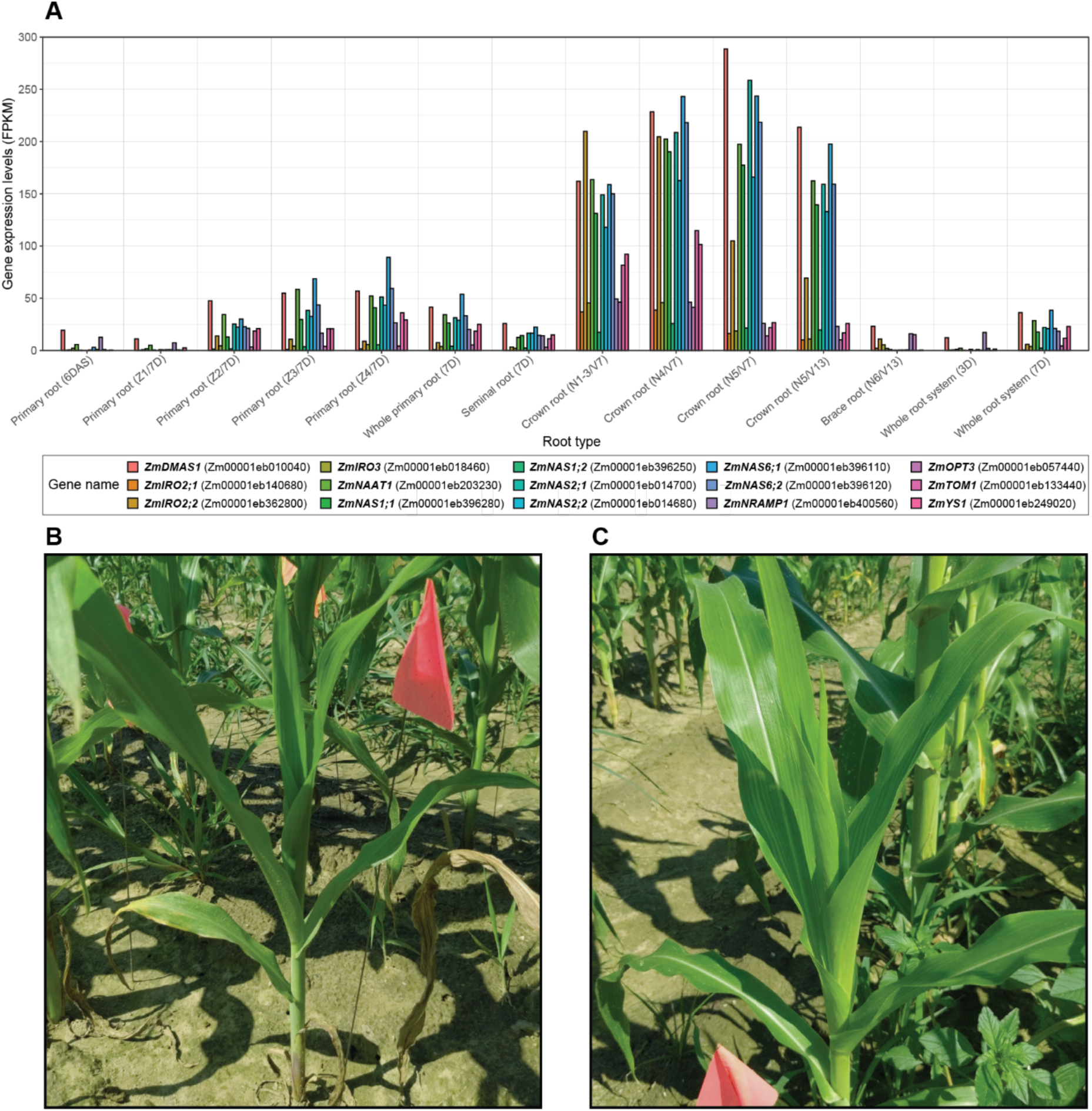
(A) Expression of 15 iron-regulated genes in different root types and developmental stages. The expression levels are given in FPKM and were extracted from the Stelpflug et al 2016 dataset (Stelpflug et al., 2016). (B) Above-ground phenotype of *rt1* mutant (Hochholdinger et al., 2004) grown in the field. (C) Above-ground phenotype of *rtcs* mutant (Taramino et al., 2007) grown in the field.

**Supplemental figure S5:**
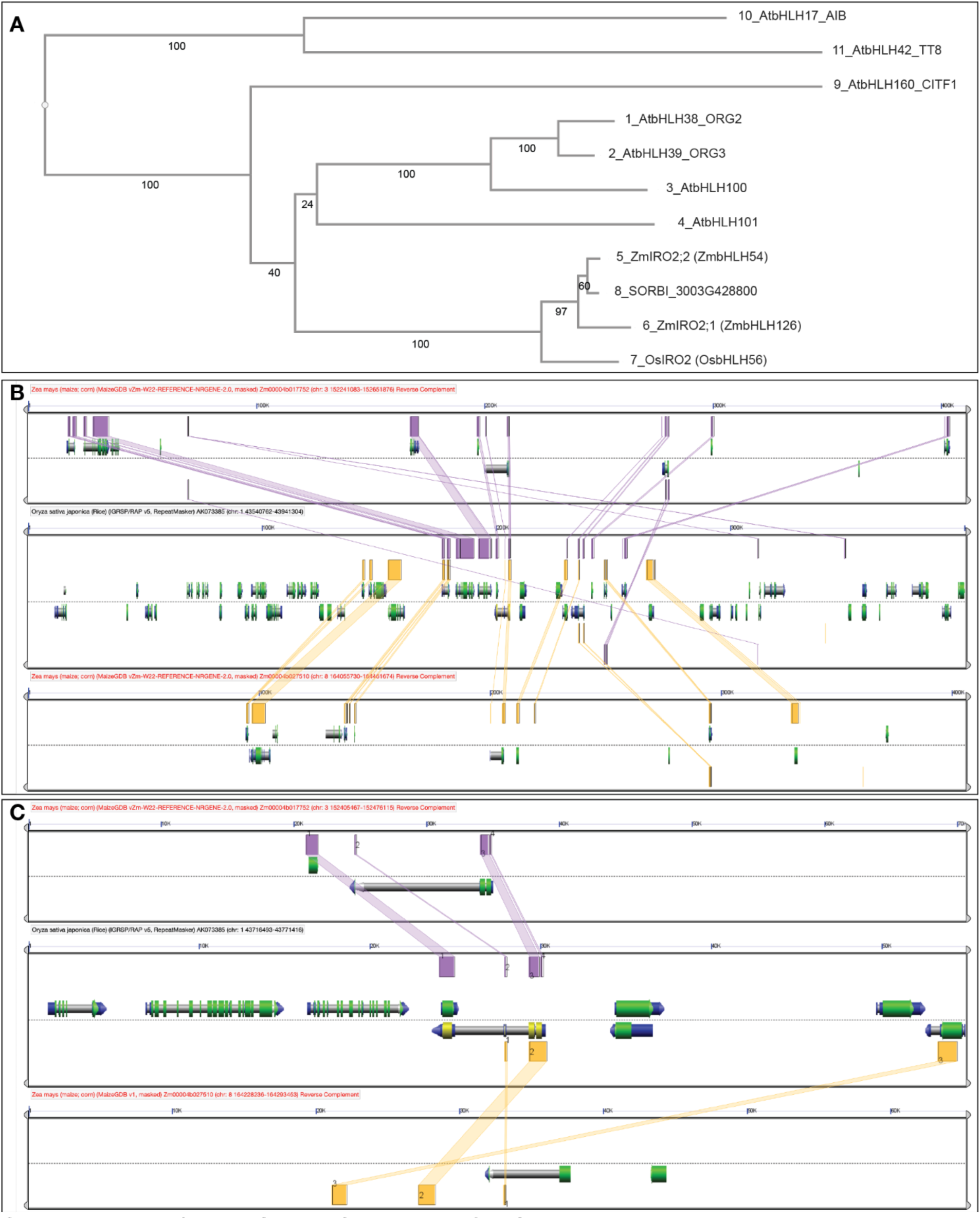
A, Gene tree of IRO2 homologs. At: *Arabidopsis thaliana*, Zm: *Zea mays*, Os: *Oryza sativa*, SORBI: *Sorghum bicolor*. B, Analysis of syntenic regions between maize and rice flanking the IRO2 genes. The central panel shows the rice chromosome 1 region, the top panel shows the maize chromosome 3 region and the bottom panel shows the maize chromosome 8 region. The gene of interest is shown in the middle of each panel. Purple connectors denote the matching regions between rice chromosome 1 and maize chromosome 3, while yellow connectors denote the matching regions between rice chromosome 1 and maize chromosome 8. C, Detailed view of B.

**Supplemental Figure S6:**
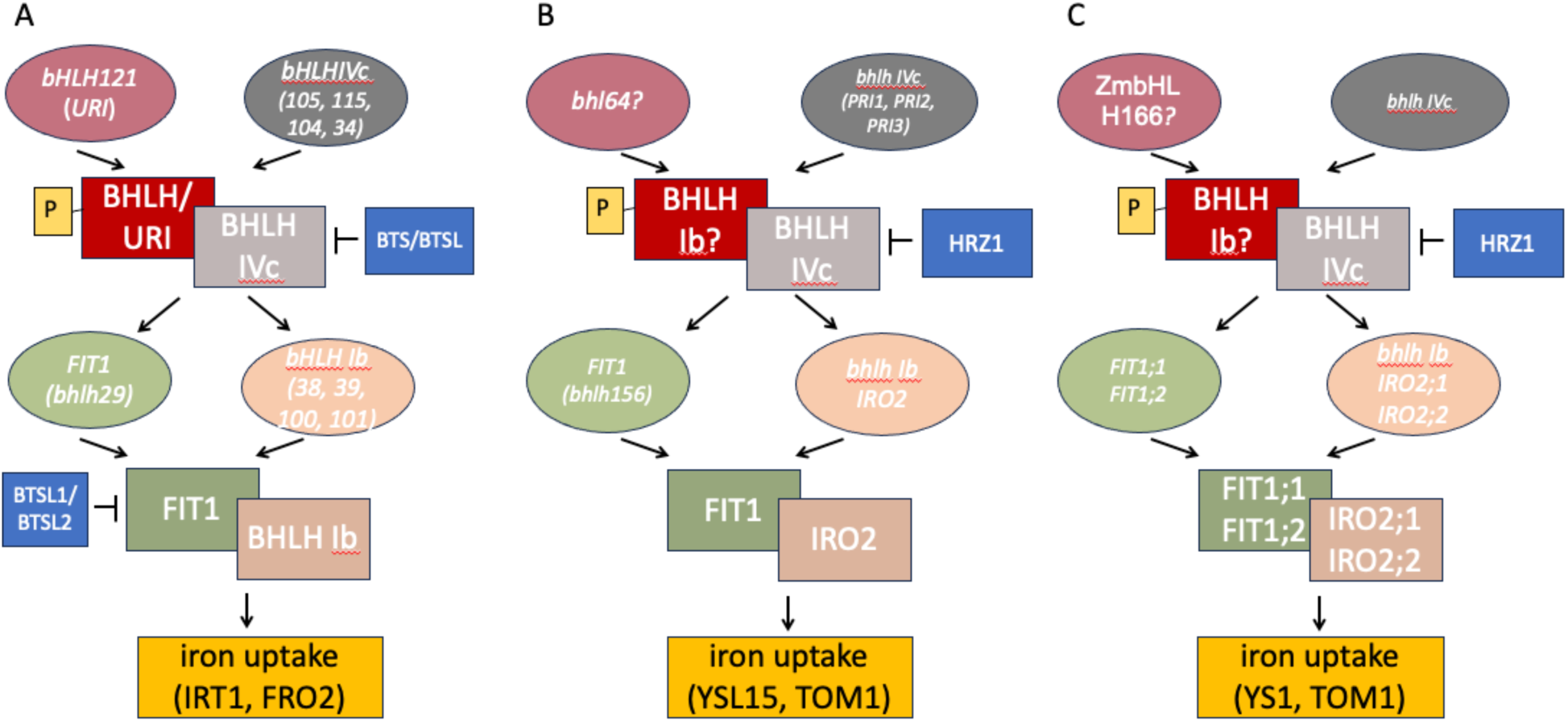
A) Highly simplified model of transcriptional control of iron uptake in Arabidopsis. BHLH121 (URI) becomes phosphorylated during iron deficiency, and the phosphorylated protein interacts with members of the bHLH IVc group (bHLH34, 103, 105, and 115) to activate expression of the next tier of bHLH genes, including FIT1 and the bhlh Ib group (bHLH38, 39, 100 and 101). FIT1 forms heterodimers with the Ib proteins, and these dimers activate genes like *IRT1* and *FRO2*, which encode core components of the iron uptake apparatus of the root. Notably, FIT1 also interacts with several other BHLH proteins to either activate or repress other aspects of iron homeostasis, including iron storage and iron transport within the plant (not show) B) Highly simplified model of transcriptional control of iron uptake in rice, highlighting parallels with the Arabidopsis mechanism. The equivalent of AtURI has not been identified in rice, but, based on sequence similarity may be OsbHLH064. The rice bHLH IVc group comprises three genes (PRI1, PRI2, and PRI3) which, possibly in combination with OsbHLH064, activate expression of OsFIT and the single rice Ib bHLH gene, IRO2. As in Arabidopsis, IRO2 and FIT interact to activate expression of genes involved in iron uptake. In the Strategy II plant, rice, the activated genes are different, though, and include genes such as YSL15 and TOM1 that are key components of grass iron uptake. C. Postulated model for transcriptional control of iron uptake in maize, highlighting parallels with Arabidopsis and rice mechanisms.

**Supplemental data 1:** Lists of iron-regulated genes generated by RNA-seq analysis of plants growing under +Fe, –Fe and ±Fe.

**Supplemental data 2:** Lists of differentially expressed genes in roots and shoots of

*ccd8* mutant seedlings.

**Supplemental data 3:** Gene ontology enrichment analysis for iron-regulated genes.

**Supplemental data 4:** List of oligonucleotide sequences used in this study.

## Notes

### Competing Interest Statement

The authors have declared no competing interest.

